# WENGAN: Efficient and high quality hybrid *de novo* assembly of human genomes

**DOI:** 10.1101/840447

**Authors:** Alex Di Genova, Elena Buena-Atienza, Stephan Ossowski, Marie-France Sagot

## Abstract

The continuous improvement of long-read sequencing technologies along with the development of ad-doc algorithms has launched a new *de novo* assembly era that promises high-quality genomes. However, it has proven difficult to use only long reads to generate accurate genome assemblies of large, repeat-rich human genomes. To date, most of the human genomes assembled from long error-prone reads add accurate short reads to further polish the consensus quality. Here, we report the development of a novel algorithm for hybrid assembly, WENGAN, and the *de novo* assembly of four human genomes using a combination of sequencing data generated on ONT PromethION, PacBio Sequel, Illumina and MGI technology. WENGAN implements efficient algorithms that exploit the sequence information of short and long reads to tackle assembly contiguity as well as consensus quality. The resulting genome assemblies have high contiguity (contig NG50:16.67-62.06 Mb), few assembly errors (contig NGA50:10.9-45.91 Mb), good consensus quality (QV:27.79-33.61), and high gene completeness (BUSCO complete: 94.6-95.1%), while consuming low computational resources (CPU hours:153-1027). In particular, the WENGAN assembly of the haploid CHM13 sample achieved a contig NG50 of 62.06 Mb (NGA50:45.91 Mb), which surpasses the contiguity of the current human reference genome (*GRCh38* contig NG50:57.88 Mb). Providing highest quality at low computational cost, WENGAN is an important step towards the democratization of the *de novo* assembly of human genomes. The WENGAN assembler is available at https://github.com/adigenova/wengan

## 1 Introduction

Genome assembly is the process by which an unknown genome sequence is constructed by detecting overlaps between a set of redundant genomic reads. Most genome assemblers represent the overlap information using different kinds of assembly graphs [1, 2]. The main idea behind these algorithms is to reduce the genome assembly problem to a path problem where the genome is reconstructed by finding “the” true genome path in a tangled assembly graph [1, 2]. The tangledness comes from the complexity that repetitive genomic regions induce in the assembly graphs [1, 2]. The first graph-based genome assemblers used overlaps of variable length to construct an overlap-graph [2]. In such graph, the reads are the vertices and the edges represent the pairwise alignments [2]. The main goal of the overlap graph approach and of its subsequent evolution, namely the string graph [2], is to preserve as much as possible the reads information [2]. However, the read-level graph construction requires an expensive all-vs-all read comparison [2]. The read-level nature implies that a path in such a graph represents a read layout, and a subsequent consensus step must be performed in order to improve the quality of bases called along the path [2]. These graph properties are the foundation of the overlap-layout-consensus (OLC) paradigm [2–4].

A seemingly counterintuitive idea is to fix the overlap length to a given size (*k*) to build a *de Bruijn* graph [1]. However, *de Bruijn* graphs have several favorable properties making them the method of choice in many modern short-read assemblers [5–7]. In this approach, the fixed-length exact overlaps are detected by breaking the reads into consecutive *k*-mers [1]. The *k*-mers are usually stored in hash tables (constant query time), thus avoiding entirely the costly all-vs-all read comparison [5–7]. A *de Bruijn* graph can then be constructed using the distinct *k*-mers as vertices, adding an edge whenever two vertices share a *k* − 1-overlap. Unlike a string graph, the *de Bruijn* graph is a base-level graph [1, 5–7], thus a path (contig) represents a consensus sequence derived from a pileup of the reads generating the *k*-mers (*k*-mer frequency). Moreover, the *de Bruijn* graph is useful for characterizing repeated as well as unique sequences of a genome (repeat graph [1]). However, by splitting the reads into *k*-mers, valuable information from the reads may be lost, especially when these are much longer than the selected *k*-mer size [2].

The type of overlap detected, and therefore the type of assembly graph constructed is intrinsically related to the sequencing technology used to generate the reads. Modern high-throughput sequencing machines are divided in two classes. One class produces short (100-300 bp) and accurate (base-error <0.1%) reads [8, 9] and a second class produces long (>10kb) but error-prone (base-error < 15%) reads [10, 11]. Despite the high per base error rate of long-reads, the latter are the better choice for genome reconstruction [12], as longer overlaps reduce the complexity of the assembly graph [13], and therefore more contiguous genome reconstructions are achievable [12]. Independent of the sequencing technology, the goals of a genome assembler are to reconstruct the complete genome in (1) the fewest possible consecutive pieces (hopefully chromosomes) with (2) the highest base accuracy while (3) minimizing the computational resources (the 1-2-3 Goals). Clearly, short-read *de Bruijn* graph assemblers are good for accomplishing goals 2 and 3 [5–7], while long-read assemblers excel at achieving goal 1 [3, 4].

Modern long-read assemblers widely adopted the OLC-paradigm [3, 4, 14–17] and new algorithms have substantially accelerated the all-vs-all read comparison [14–17]. Such progress has been possible by avoiding entirely the long-reads error-correction step [14–17], and by representing the long-reads as fingerprints derived from a subset of special *k*-mers (*i.e.* minimizers [18], homopolymers compressed *k*-mers [14], minhash [17] etc.). The reduced long-read representation is appropriate for detecting overlaps >2kb in a fast way [14, 16, 17]. The newest long-read assemblers are therefore starting to be good also at goal 3 [14, 16, 17]. However, assembling uncorrected long-reads has the undesirable effect of giving more work to the consensus polisher [15,17,19–21]. Genome assembly polishing is the process of improving the base accuracy of the assembled contig sequences [15, 17, 19–22]. It can be applied during and after the genome assembly. Usually, long-read assemblers perform a single round of long-read polishing [14,16,17], that is followed by several rounds of polishing with long [15, 17, 19, 21] and short [15, 20, 22] reads using third-party tools [15, 17, 19–22].

Currently, polishing large genomes, such as the human genome, can take much more computational time than the long-read assembly itself [14, 16, 17]. Moreover, there is no standard practice for polishing large genomes, and usually several rounds of polishing are employed with user-defined criteria in order to remove consensus errors, notably the short-indels occurring in homopolymer regions, which are the characteristic error signature of current long-read technologies [14, 16, 17]. This new assembly approach gave rise to some criticisms because even after several rounds of polishing, a substantial fraction of consensus errors remains, hampering the subsequent genome analyses such as gene and protein prediction [23].

When the aforementioned assembly approach employs short-read polishing [15, 20, 22], then it corresponds to a long-read-first hybrid assembly strategy [24, 25]. Another hybrid assembly strategy consists in starting the assembly process with short reads [26]. One short-read-first implementation scaling to human size genomes starts with a *de Bruijn* graph approach, followed by error-correction of long-read sequences using the short-read-contigs, and finally assembly of corrected long-read sequences using the OLC paradigm [26]. Notice that none of the described hybrid strategies employs the short-reads to tackle the problem of assembly contiguity (*i.e.* they do not aim at joining two long reads by a short-read contig). They therefore exploit only partially the short-read sequence information.

In this paper, we introduce WENGAN, a new hybrid genome assembler that unlike most state-of-the-art long-read assemblers, entirely avoids the all-vs-all read comparison, does not follow the OLC paradigm, and integrates short-reads in the early phases of the assembly process (short-read-first). WENGAN implements efficient algorithms that exploit the information of short and long reads to tackle contiguity as well as consensus accuracy. Thus, WENGAN is a full hybrid assembler. We validated WENGAN with standard assembly benchmarks. Our results demonstrate that WENGAN is the only genome assembler that optimizes the 1-2-3 Goals and is particularly effective at low long-read coverage (15X). Furthermore, we show that the WENGAN assemblies performed by combining short and ultra-long Nanopore reads surpass the contiguousness of the current human reference genome.

## 2 Results

### 2.1 The WENGAN algorithm

Accurate short-reads are good for assembling unique genomic regions and bad for solving repeats. Long-reads are good for assembling any genomic region, but bad at consensus generation due to their high error rate (15%). Thus, a good hybrid assembler can overcome the weaknesses of both technologies by exploiting their complementary features (accuracy and length). We describe here the main steps of WENGAN and its key algorithmic ideas for enabling efficient hybrid assemblies of human genomes (for more details see the Methods section).

WENGAN starts by building short-read contigs using a *de Bruijn* graph assembler [5–7] (Figure 1A1). Then, the pair-end reads are pseudo-aligned [27] back (pseudo because we use an alignment-free approach for this) to detect and error-correct chimeric contigs as well as to classify them as repeats or unique sequences (Figure 1A2). Repeated sequences induce complex *de Bruijn* graph topologies in their neighborhood, and short-read assemblers can choose wrong paths while traversing such complex regions, thus leading to chimeric contigs (Figure S1). Chimeric short-read contigs limit the accuracy and contiguousness of the assembly when they are not corrected (Figure 1B). Each short-read contig is therefore scanned base-by-base and split at sub-regions lacking of pair-end read support (that is, lacking of physical coverage, Figure S1).

**Figure 1:**
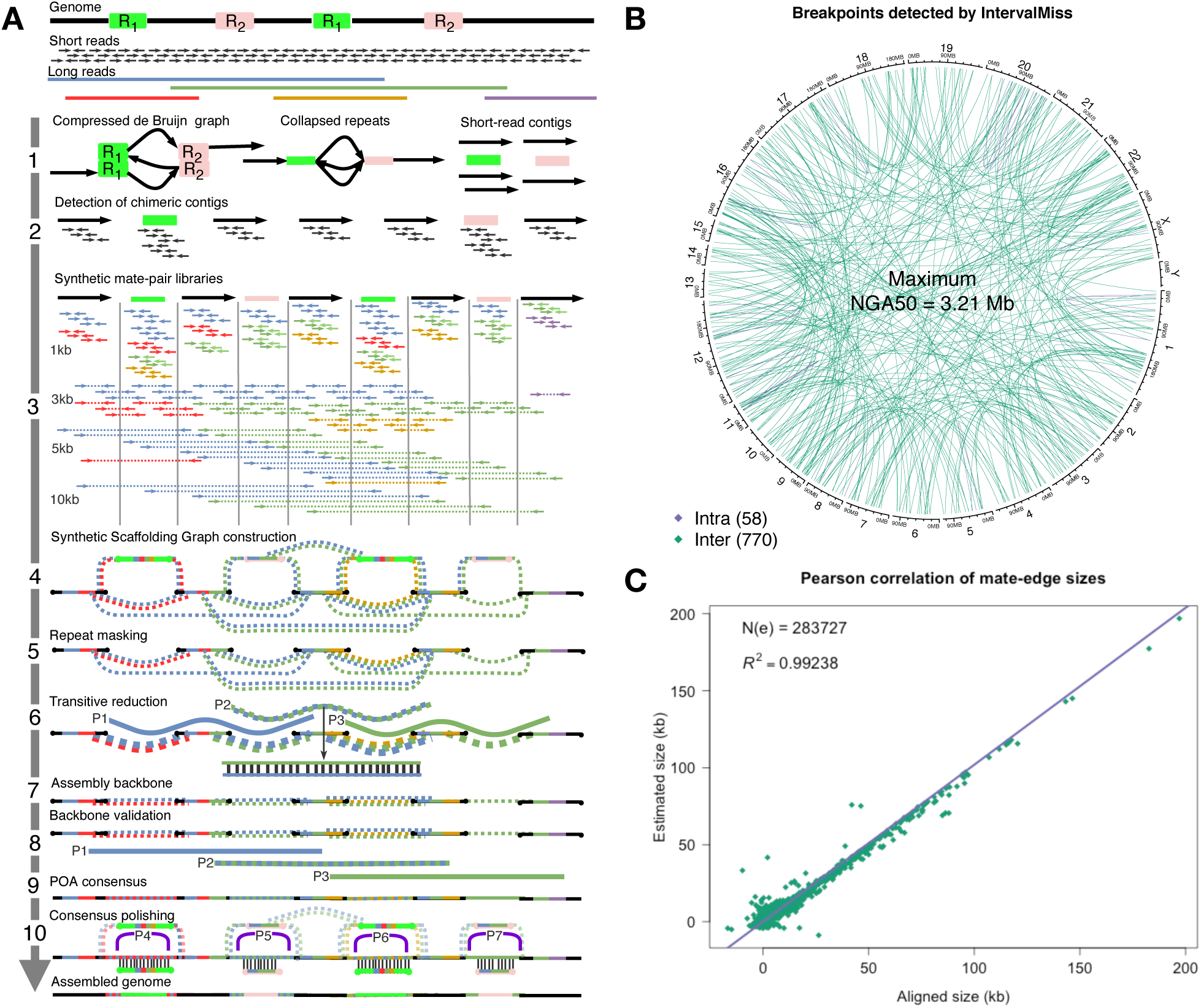
The WENGAN algorithm. The WENGAN steps consist in first assembling and error correcting the shortread contigs (1A-2A), then in creating a spectrum of synthetic mate-pair libraries from long-reads (3A) to build the Synthetic Scaffolding Graph (SSG, 4A). The SSG is used to compute approximate long-read overlaps by building long-read-coherent paths (5A-6A). The long-read overlaps restore the long-read information and facilitate the construction and validation of the assembly backbone (7A-8A). The SSG is used to fill the gaps by building for each mate-edge a consensus sequence using the Partial Order Alignment graph (9A). The final steps use the SSG to polish the consensus sequences (10A). (B) The circular plot depicts the number of missassemblies detected using the pair-end information and the maximum NGA50 that can be achieved if those contigs are not corrected in the NA12878 MINIA3 short-read assembly. (C) The Pearson correlation of the mate-edge lengths before (y-axis) and after (x-axis) building the consensus sequences for a total of 283,727 mate-edges from the NA12878 WENGAN_*M*_ assembly is depicted. Notice that the agreement between the estimated and the aligned mate-edge lengths exceeds *R*^2^ > 0.99. The repeat contigs (2A-10A) are drawn uncollapsed to explain the WENGAN steps (A).

Following short-read contigs correction, we generate synthetic paired-reads of different insert sizes from long-read sequences, which are mapped to the corrected short-read contigs (Figure 1A3). The spectrum of synthetic libraries is used to span the genomic repeats. For instance with ultra-long nanopore reads, we can create a spectrum composed of 24 synthetic libraries with insert sizes ranging from 0.5kb to 200kb (Figure S2). Each pair-end is pseudo-aligned on the fly to the corrected short-read contigs, and matched pairs are stored with a reference to the long-read from which they were extracted (colors appearing in pairs, Figure 1A3). Using the mapped pairs and the corrected short-read contigs, we then build the Synthetic Scaffolding Graph (SSG). The SSG is an extension of the scaffolding graph [28], where there is an additional edge-labeling function that labels (colors) the SSG edges with the long-reads. The labeling is done by using as a proxy the pseudo-aligned synthetic pairs (Figure 1A3-4). After the SSG construction (Figure 1A4) and subsequent repeat masking (Figure 1A5), we employ the SSG to compute implicit approximate long-read multiple alignments by searching for alternative long-read-coherent paths (Figure 1A6). The aim of this graph operation (called *transitive reduction*) is to restore the full long-read information in the SSG. Each successful reduction modifies the weight as well as the shape of the SSG (Figure 1A6).

After restoring the long-read information, we order and orient the short-read contigs by applying an approximation algorithm [29] that uses all the connectivity information at once to produce an optimal assembly backbone (lines or scaffolds, Figure 1A7). The solution is validated by checking the distance constraints that the reduced long-read-coherent paths impose on the assembly backbone (physical long-read-coherent paths coverage, Figure 1A8).

A property of the SSG is that all the edges connecting two short-read contigs (called *mate-edges*) are spanned by at least one long-read. We therefore use the inner long-read sequence of the synthetic mate-pairs that span the mate-edge to build a long-read consensus sequence using a partial order alignment graph [15,30] (Figure 1A9). The corresponding short-read contig-ends are then aligned [31] to the mate-edge consensus sequence to determine the correct boundaries, thus filling the gap between the two short-read contigs. We computed the Pearson correlation of the mate-edge length before and after filling the gap for a total of 283,727 mate-edges. The correlation is very high (R^2^ > 0.99) even for large gaps (>100kb, Figure 1C).

The final steps use the SSG to polish the mate-edge consensus sequences by finding long-read-coherent paths that traverse the repeated regions (*i.e.* P4 and P6 for R1, Figure 1A10) or pairwise alignments [31] between the repetitive short-read contigs and the mate-edge consensus sequences (Figure 1A10). Finally, the hybrid contigs are reported in FASTA format (Figure 1A).

### 2.2 WENGAN optimizes the 1-2-3 *de novo* assembly goals

To validate WENGAN, we used public Oxford Nanopore reads [24] (rel5) and Illumina short-reads (Table S1) of the NA12878 human cell line. In particular, rel5 has a total of 11.6 million reads (40X) with an N50 of 13.6kb (that is, 50% of the total sequence data is in reads longer than 13.6 kb), and 3.3X of the genome is covered by reads longer than 100kb (Table S2). WENGAN was bench-marked in its three assembly modes, namely WENGANM (MINIA3 [5]), WENGANA (ABYSS2 [6]) and WENGAND (DISCOVARDENOVO [7]). WENGAN modes reflect the short-read assembler used (Figure 1A1, Methods section) and are intended to adapt WENGAN to different computational resources. We compared WENGAN to four state-of-the-art assemblers, MASURCA (hybrid short-read-first) [26], CANU (long-read only) [4], WTDBG2 (long-read only) [14], and FLYE (long-read only) [16]. All genome assemblies, which we compared in this benchmark were generated by the developer of the respective assembler (Table S3). The benchmark is described in detail in the Methods section (Assembly validation).

WENGAN produced the most contiguous assemblies, with contig NG50s of 16.67, 23.08, and 33.13 Mb, for WENGANM, WENGANA and WENGAND, respectively (Table 1). The best long-read assembler among the four evaluated, namely FLYE (NG50 22.18Mb), is comparable to WENGANA (NG50 23.3Mb), but is surpassed by WENGAND (NG50 33.3Mb). All the other evaluated assemblers are outperformed by any WENGAN mode (NG50 >= 16.67Mb, Table 1). Moreover, WENGAN increased the contiguousness of the short-read-only assemblies by a factor of 1722X, 1783X and 362X, for MINIA3 (NG50 9.6kb), ABYSS2 (NG50 12.9kb) and DISCOVARDENOVO (NG50 91kb), respectively (Table S5).

**Table 1:**
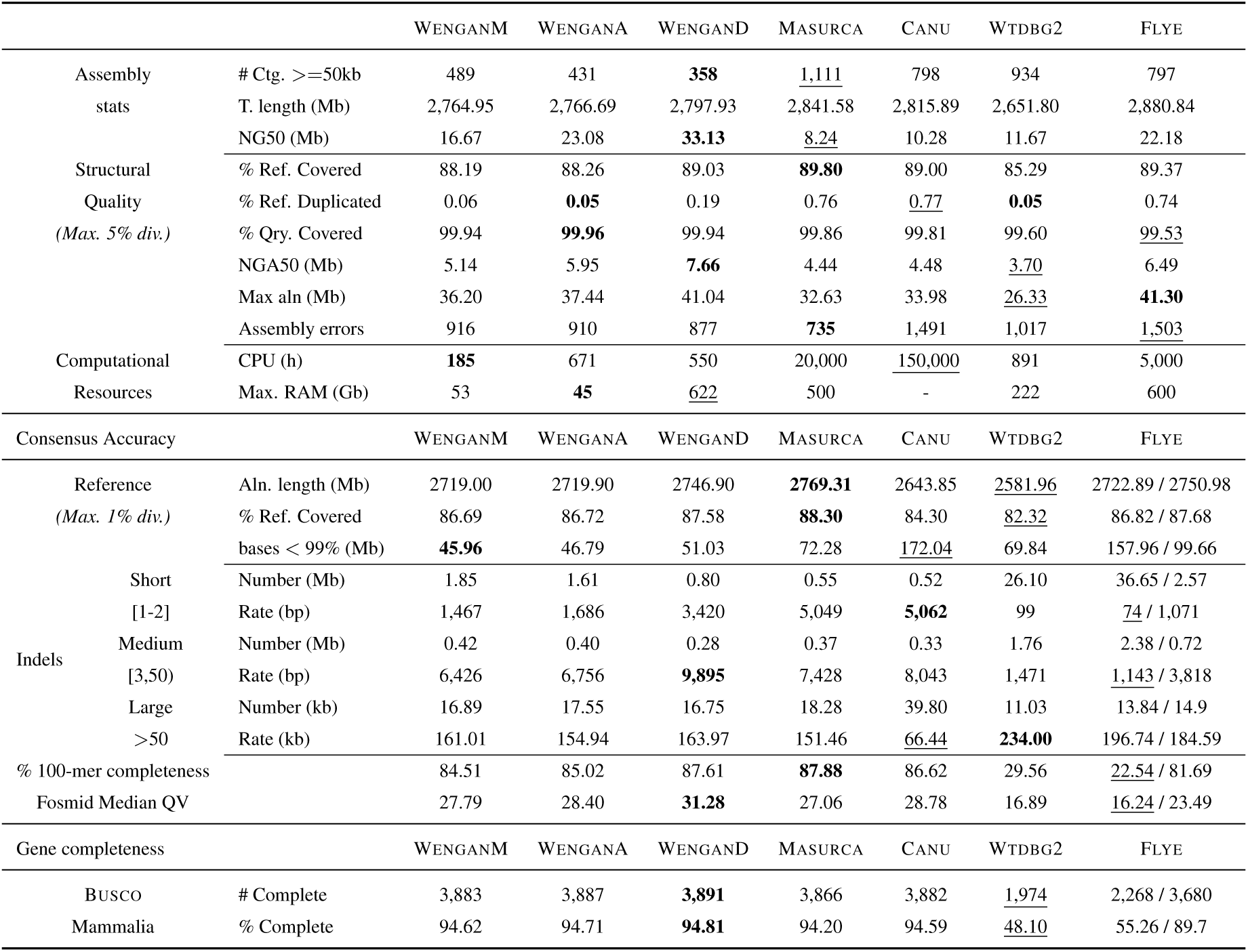
Quality assessment of the *de novo* assemblies of NA12878. Structural and consensus accuracy was determined as described in detail in the Methods section (Assembly validation). All the NA12878 assemblies were built by the assembler developpers using the Oxford Nanopore (rel5) plus Illumina data at the assembly or polishing steps (Except WTDBG2). The CANU assembly was hybrid polished with NANOPOLISH × 2, RACON × 2, and PILON × 2. The FLYE assembly was hybrid polished with RACON × 2 and NTEDIT × 3 (Method section). The WENGAN, MASURCA and WTDBG2 assemblies were not polished by external tools. The reported CPU time does not include the CPU time spent polishing the assembly with external tools. NG50 is the contig length such that using longer contigs produces half (50%) of the bases of the reference genome (3.1374 Gb). NGA50 is NG50 corrected of assembly errors. The sentence “bases < 99%” in the table corresponds to the number of assembly bases that cannot be aligned in blocks ≥ 50kb at average identity ≥ 99%. The variant-rate was computed dividing the amount of assembly sequence aligned by the number of variants called on such alignments. The 100-mers completeness is the fraction of distinct 100-mers in the reference (2.825 Gb) that are captured in the corresponding assembly. Consensus statistics before and after the polishing are included for the FLYE assembly.

The structural quality was determined by whole genome alignment of the assembled contigs against the human reference genome (see Assembly validation in the Methods section). Aligned blocks larger than 50kb with an average identity ≥ 95% were used to collect the metrics (Table 1). The WENGAN assemblies cover more than 88% of the reference with more than 99.9% of the assembled sequence mapped, and the contigs have fewer duplicates than the contigs of its competitors (except WTDBG2, Table 1). The NGA50 (which corresponds to the NG50 corrected of assembly errors) of WENGAN is better than the one of MASURCA, WTDBG2 and CANU (Table 1). FLYE has an NGA50 that is better than WENGANM, is comparable to WENGANA and is lower than WENGAND (Table 1). A QUAST [32] analysis with reduced alignment identity (≥80%) which favors long-read assemblers, reveals that WENGAN (10.9Mb-14.2Mb) has a better NGA50 than all its competitors and almost double the NGA50 of MASURCA (5.2Mb), WTDBG2 (6.2Mb) and CANU (6.4Mb) (Table S6). Moreover, WENGAN consistently showed a lower number of assembly errors than its peers (Table 1, Table S6).

The consensus accuracy was determined using different sequence analyses (see Assembly validation in Methods section). The level of polishing of the evaluated assemblies goes from none to complete. In particular, assemblies generated by WENGAN, MASURCA or WTDBG2 were not polished by external tools. The CANU assembly was polished with short and long reads (2 x NANOPOLISH [21], 2 × RACON [15] and 2 × PILON [20]). Similarly, we polished the FLYE assembly with short and long reads (2 × RACON [15] + 3 × NTEDIT [22], see Methods). All assemblies were aligned to the reference genome and the aligned blocks ≥ 50kb with an average identity ≥ 99% were used to call variants (Table 1). The WENGAN assemblies cover ≥ 86.6% of the reference, which is better or comparable to the other assemblers, and has the lowest amount of unaligned assembled sequences at 99% identity (Max. 51.03Mb, Table 1). We computed the rate of short(1-2), medium(3-50] and long(≥50) indels for each of the genome assemblies (Table 1). CANU-POLISHED and MASURCA have better short indel rates than the WENGAN assemblies, but WENGAN has better than or comparable medium and long indel rates (Table 1). Moreover, unlike long-read assemblers, the majority (≥72%) of the WENGAN consensus errors are located in the mate-edge consensus sequences (Figure S3). The 100-mer analysis reveals that the WENGAN assemblies contain more than 84.5% of the 100-mers of the reference (Table 1). The alignment of 103 curated random Fosmid sequences of NA12878 [7] (3.92Mb, Table S7) allows us to estimate that the median consensus quality of the WENGAN assemblies is QV ≥ 27.79 (Table 1, maximally one error every 625 bases). The median QV values of WENGAN are better or comparable to the values of CANU-POLISHED and MASURCA (Table 1). The BUSCO gene completeness of WENGAN assemblies ranges from 94.62% to 94.81%, which is higher than the result of any other evaluated assembler and reflects the good consensus quality and contiguity of WENGAN‘s assemblies (Table 1).

FLYE and WTDBG2 consistently were the worst in the consensus quality benchmark because of the poor polishing performed. Moreover, polishing the FLYE assembly with long reads and with the same short reads used by WENGAN consumed 755 CPUs hours (Table S8). While the hybrid polishing removed millions of consensus errors (Table 1, Table S8), and increased the median quality value and the BUSCO gene completeness (to 23.39 and 89.7%), the polished FLYE assembly still has a lower quality than any of the unpolished WENGAN assemblies (Table 1).

In terms of computational resources, the WENGAN assemblies consumed less CPU hours than any other method (Table 1). WENGANM, the fastest WENGAN mode based on MINIA3, consumed 810 times less CPU-hours than CANU (185 vs.∼150,000 hours, Table 1) and only required 53Gb of RAM to complete the assembly (Table 1).

Collectively, the benchmark results demonstrate that WENGAN is the only genome assembler evaluated that optimizes all of the 1-2-3 *de novo* assembly goals, namely, contiguity, consensus accuracy, and computational resources.

### 2.3 WENGAN is effective at low long-read coverage

In order to optimize the cost of generating high quality genome assemblies, we next investigated the required long-read coverage to produce *de novo* assemblies with NG50 of at least 10Mb. Moreover, we assessed the suitability of the BGI sequencing technology [9] (MGIseq-2000) as an alternative to Illumina SBS [8] for hybrid assembly using matched short-read genomic data. We sequenced the NA12878 human cell line using the short-read sequencers NovaSeq-6000 [8] and MGIseq-2000 [9] as well as the long-read sequencer ONT PromethION [11] (Methods section). We generated a total of 548.2 million pair-end reads (2×150bp) of sequence (53.06X) from both short-read sequencers (Table S1, Method section). Futhermore, three flow-cells of PromethION produced a total of 10.4 million reads (40X) with a N50 of 17.18 (kb) (Table S2, Methods section). The Nanopore reads were base-called using GUPPY (v3.0.3) with the high accuracy FLIP-FLOP model. We randomly subsampled the long-read data from 10X to 30X of genome coverage in increasing batches of 5X. The N50 was nearly identical for all the long-read subsamples (N50 = 19.6 kb, Table S9). The number of pair-end reads was fixed to 548.2 million for both short-read technologies (Table S2). WENGAN and the best long-read assembler among those evaluated, namely FLYE (v2.5), were used to build hybrid and long-read assemblies for each subsample (Figure 2, Table S10).

**Figure 2:**
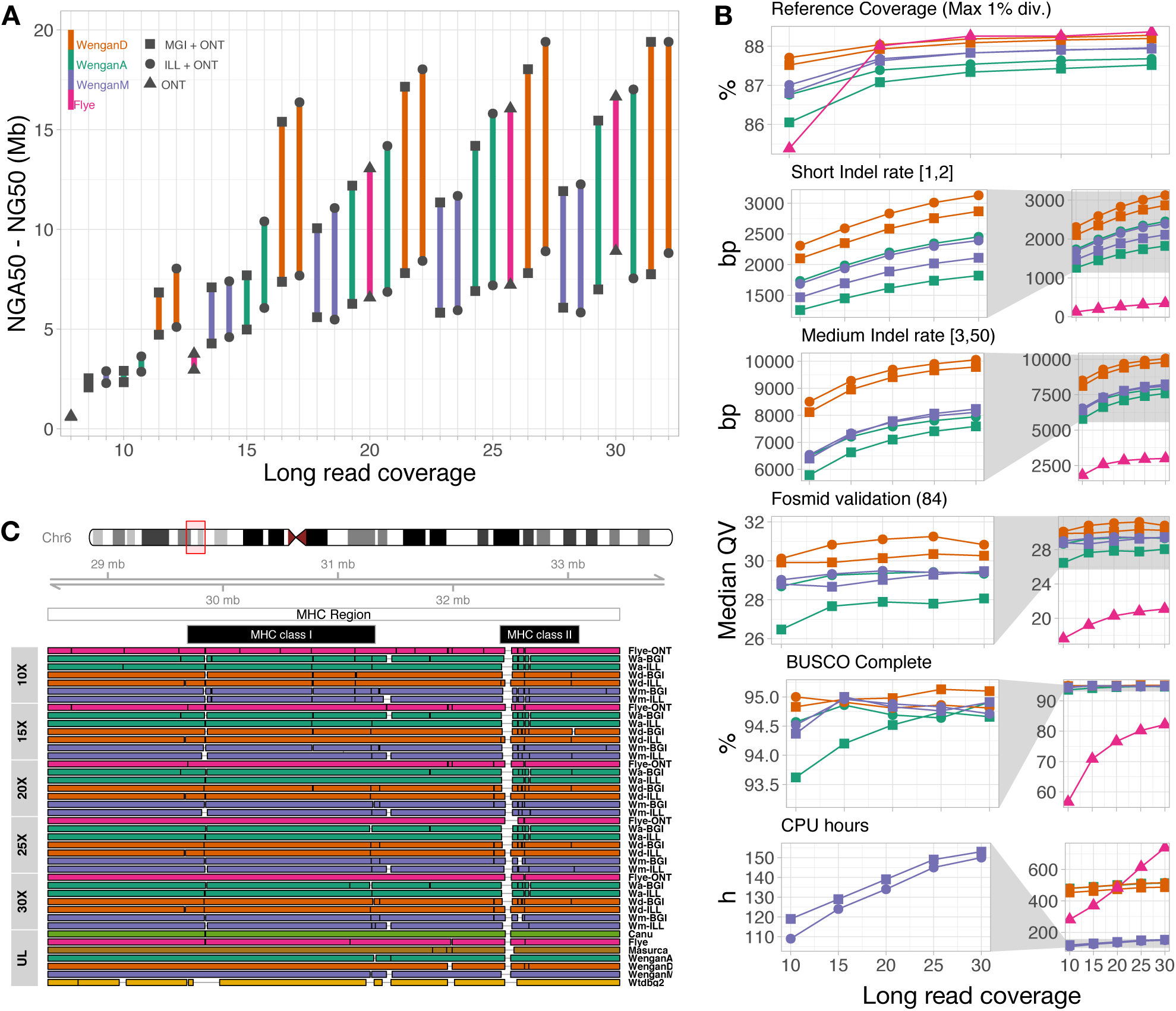
*De novo* genome assemblies of NA12878 varying the long-read coverage and the short-read technology. A) The *de novo* assemblies were sorted by NG50 at each long-read coverage (lolliplot). We computed the NGA50 (which corresponds to the NG50 corrected of assembly errors) of each assembly by a whole genome alignment against the reference allowing a maximum of 5% divergence (see Methods section). B) The consensus quality (see Methods section) of each genome assembly as well as the CPU-hours required for the assembly are reported. C) The WENGAN and FLYE assemblies of the complex MHC region located in Chr6:28,477,797-33,448,354 (4.97Mb). The MHC sequence was aligned to the genome assemblies and the aligned blocks ≥ 30kb with a minimum identity of 95% were kept. The alignment breakpoints (vertical black lines) indicate a contig switch, alignment error or gap in the assembly. The assemblies of the MHC region are displayed in tracks by long-read coverage. The track UL shows the assemblies of the MHC region using the ultra-long nanopore reads (rel5).

A major increase in contiguity for WENGAN was observed when going from 10X to 15X of long-read coverage (Figure 2A). In particular, we observed an NG50 increase from 2.5, 2.9, 6.8 Mb to 7.1, 7.7, 15.4 Mb for WENGANM, WENGANA, and WENGAND, respectively. The major gain in contiguity for FLYE occurred at 20X when its NG50 increases from 3.7 (15X) to 13.02Mb (Figure 2, Table S10). At shallow long-read coverage (10-15X), FLYE is outperformed by all WENGAN modes. Over 20X of coverage, FLYE outperforms WENGANM and is comparable in contiguity to WENGANA (Figure 2). Notably, WENGAND using 15X of long-read coverage leads to an NG50 of 15Mb, which FLYE can only reach at 30X of long-read coverage (Figure 2A). All WENGAN modes at 20X of coverage produced hybrid assemblies with NG50 ≥ 10Mb and NGA50 ≥ 5.4Mb (Figure 2A, Table S10).

All assemblies generated by WENGAN cover more than 86.5% of the reference genome at any long-read coverage (Max 1% divergence, Figure 2B). As expected, FLYE achieves its highest consensus quality at 30X of long-read coverage (max QV=21.08, Figure 2B). Polishing FLYE with long and short (NovaSeq) reads increased its median consensus quality to QV=27.21 (Table S12). Almost all WENGAN assemblies achieve higher consensus quality than the polished FLYE assembly (except the WENGANA-BGI-10X with QV=26.47), as showed by the different sequence analyses performed (min WENGAN QV=27.67 excluding WENGANA-BGI-10X, Figure 2B, Table S10 and Table S11).

The contiguity and consensus quality of WENGAN assemblies vary more as a function of the WENGAN‘s mode than with the type of short-read data used (Figure 2A-B). Indeed, under the same WENGAN mode, the largest difference in contiguity between the short-read technologies of Illumina and BGI is NG50=2.7Mb (WENGANA at 15X, Figure 2A) and their consensus quality is almost identical (Figure 2B). WENGANM required a maximum of 153 CPU hours (max elapsed time <16.7 hours on 20 CPUs) and 44 Gb of RAM to complete the assemblies (Figure 2B, Table S10). To our knowledge, this is the first time that a genome assembler reaches a contiguity of 10Mb and consensus quality of QV 29.4 on such minimal and accessible computing resources.

The Major Histocompatibility Complex (MHC) region is hard to assemble due to its repetitive and highly polymorphic composition [24]. We checked the assemblies of FLYE and WENGAN to see if they solved the 4.97Mb MHC region (Figure 2C). The WENGAN assemblies at low coverage (≤ 20X) reach higher NGA50 than the FLYE assemblies (Figure 2C, Figure S4). However, FLYE over 25X of coverage assembles the MHC region in less than two contigs with a NGA50 of 4Mb (Figure 2C, Figure S4). WENGAN with ultra-long reads achieves NGA50s of 2.8Mb and spans the region with less than 4 contigs (Figure 2C-track-UL, Figure S4). The evaluated assemblers produce a mix of haplotypes of the MHC region. The assembled sequence therefore does not represent valid haplogroups and subsequent phasing must be performed to solve the region [24].

In summary, we have demonstrated that the MGIseq-2000 platform is comparable in all the assembly metrics to the Illumina NovaSeq-6000 platform, while being more cost-effective (approx. 50% lower per-base price). In addition, we demonstrated that WENGAN reduces the computational and the long-read sequencing cost of assembling a human genome. Notably, WENGAN produced a high quality assembly with NG50 > 10Mb (QV > 29) for a price of less than $1500 by combining 20X long-read coverage generated on ONT PromethION with 50X short-read coverage generated on MGI-seq2000, using less than one day of computing time on an average server (20 cores, ≤50Gb RAM). Hence, we have demonstrated that genome *de novo* assembly is now highly affordable and readily accessible to a large range of research groups.

### 2.4 WENGAN surpasses the contiguousness of GRCh38

To explore the contiguousness limit of WENGAN, we assembled the genomes of two diploid human samples, HG00733 and NA24385, and of one haploid human sample, CHM13, which were sequenced with very long reads (Table S2). All sequencing data were obtained from public repositories (Table S1 and Table S2). In particular, HG00733 was sequenced using the Pacific Biosciences technology (Sequel I) at 90X of genome coverage with N50 ≥ 33.2kb (Table S2). NA24385 and CHM13 were sequenced using the Oxford Nanopore technology at 60X and 49X of genome coverage and N50s of 54kb and 72kb, respectively (Table S2). The sequence data of NA24385 and CHM13 were generated using an ultra-long-read protocol [24] for ONT MinION and contain about 15X genome coverage of reads larger than 100 kb (Table S2). Those reads are useful to tackle the most complex human genome repeats such as large segmental duplications that often result in contig breaks in current long-read assemblies [33].

The WENGAND assembly of HG00733 has the fewest gaps of any PacBio assembly of a human genome, with more than half of the genome contained in contig sequences at least 33 Mb long (Figure 3A), a substantial improvement in accuracy and contiguity over the FALCON assembler developed by PacBio (Figure 3 and Figure S5). However, the WENGAND assembly does not surpass the contiguity of the human reference due to the lack of ultra-long-reads (Table S2). Using ultra-long-reads as input, WENGAN created a spectrum of synthetic mate-pair libraries in the range of 0.5kb-500kb for the assemblies of NA24385 and CHM13. The WENGAND assembly of NA24385 surpasses the contiguity of the GRCh37 (patch 13) reference and is just 5.8Mb less contiguous than the GRCh38 (patch 19) reference (Figure 3A and Figure S5). The WENGAND assembly almost doubles the contiguousness of the NA24385 assembly done with PacBio HiFi reads [34](CANU max NG50=27.4Mb). To our knowledge, this is the most contiguous genome assembly of a diploid human sample ever made, although the assembly has to be phased to represent properly paternal and maternal haplotypes [34].

**Figure 3:**
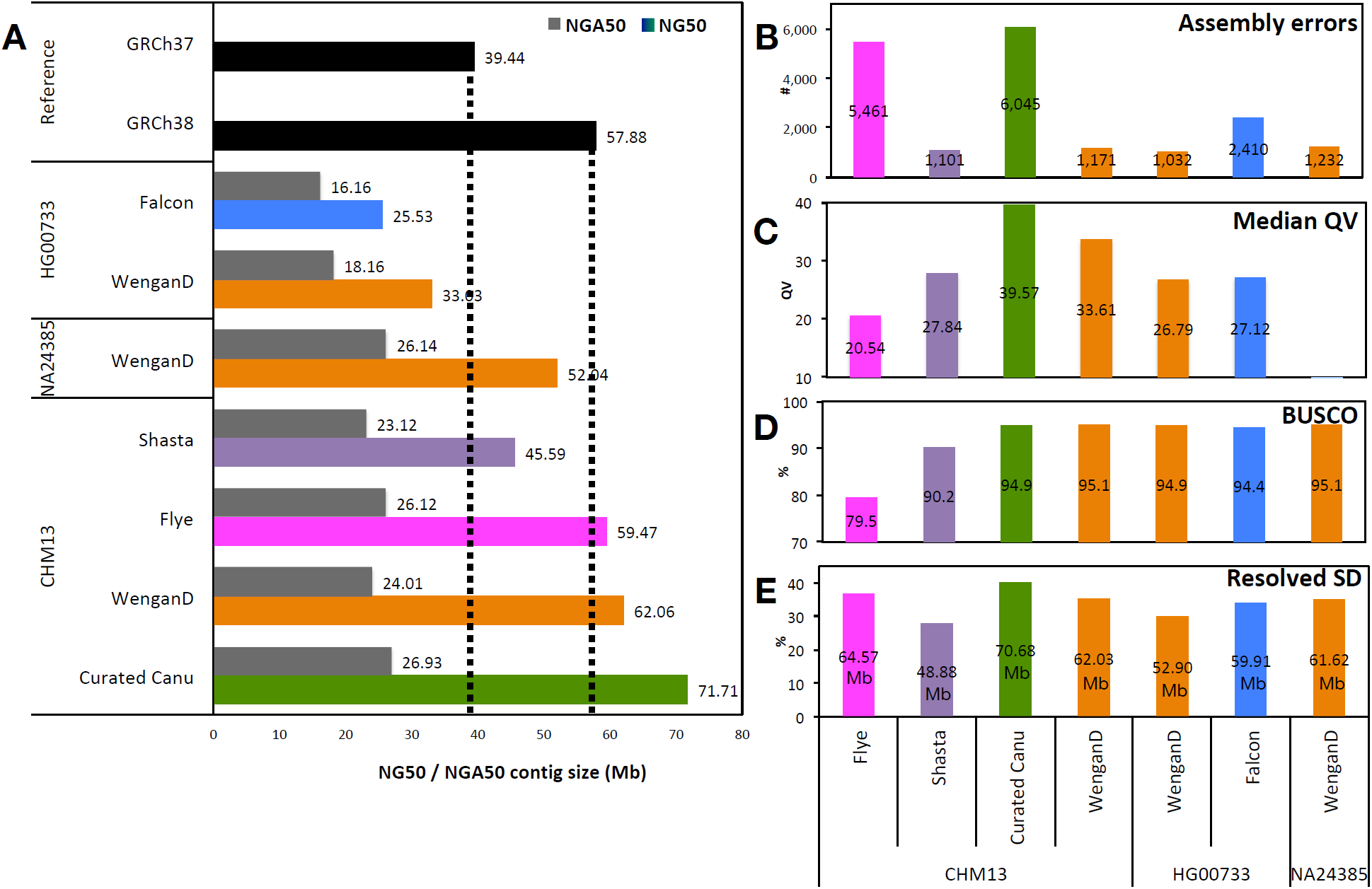
WENGAND assembly results on the CHM13, HG00073, and NA243875 genomes. A) The barplot shows the contig NG50/NGA50 of WENGAND, of the other long-read assemblers and of the current human reference genomes. NG50 and NGA50 were computed using as genome size the total contig lengths of GRCh38 (2.94Gb). B) Assembly errors predicted by QUAST using as reference GRCh38 (autosomes + X and Y). C) Consensus quality assessment by alignment of BAC sequences to the assembled contigs using the BACVALIDATION tool. D) The gene completeness was determined using the BUSCO tool. E) Segmental Duplications (SD) resolved by the genome assemblies. An SD is considered resolved if the aligned contig extends the SD flanking sequences by at least 50kb (See the Methods section). The curated CANU assembly (V0.6) is not directly comparable to the other assemblies of CHM13 because it was corrected/curated using linked-reads (10X) and Optical Mapping (BioNano) data and the centromere of chromosome X was manually resolved [25].

The WENGAND assembly of CHM13 has a total length of 2.84Gb with half of the genome contained in contig sequences larger than 62.06 Mb (NG50, Figure 3A). The contig NG50 of the WENGAND assembly surpasses the contiguousness of both human reference genomes (Figure 3A and Figure S5). We estimate the median consensus quality of this WENGAND assembly to be ≥ 99.96% (QV=33.61, Figure 3C, Table S14). Additionally, the complex MHC region is spanned in a single contig on this assembly due to the haploid nature of CHM13 sample (NGA50 2.79 Mb, Figure S6).

Consistent with our comparisons to long-read assemblers, the WENGAND assemblies of these three human samples exhibit a lower mis-assembly count (avg. 1,145) and the highest BUSCO gene completeness (≥ 94.9%, Figure 3BD). Moreover, the WENGAND assembly of CHM13 took 1,027 CPU hours (40 hours real-time on 44 CPUs) and used a maximum of 647Gb of RAM (Table S13). Such performance is at least 213X more efficient than the CANU performance [17] (*∼*219,000 hours) and used lower memory than the FLYE (v.2.5) and SHASTA (nanopore-read only) [17] assemblers.

We compared the amount of segmental duplications (SDs) resolved in the assemblies by checking the span of the 175Mb of sequence annotated as SD in GRCh38 [33]. The most contiguous WENGAN assemblies resolve over 61.5Mb of the SDs, which is better than SHASTA, comparable to FLYE and lower than the curated CANU assembly [25] (Figure 3E). With PacBio reads, the FALCON assembler resolves 7Mb more SD sequence than WENGAND (Figure 3E). However, none of the assemblers evaluated resolved more than 40% of such hard-to-assemble regions (Figure 3E). Even with ultra-long-reads further improvement of algorithmic approaches will be necessary to complete the assembly of SDs [33].

### 2.5 Evaluation of assembly accuracy and contiguity using BIONANO optical mapping

We observed that the distance between the NG50 and NGA50 values increases at greater assembly contiguousness (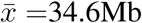, Figure 3A), which is likely caused by real sequence variation between the sequenced CHM13 sample and the GRCh38 reference genome. Given this limitation of the reference-based validation, we additionally used an independent *de novo* BIONANO genome map of CHM13 [25] to assess the correctness of the WENGAND assembly. The BIONANO map is 2.97 Gb in size with 255 contigs and an N50 of 59.6 Mbp. The BIONANO map is integrated with the sequence assembly by identifying *in silico* the nicking endonuclease-specific sites on the contig sequences (*in silico* map) and then both maps are aligned (Figure 4). Conflicts between the two maps are identified and resolved, and hybrid scaffolds are generated by using the Bionano maps to join the contig sequences and vice versa (Figure 4). A total of 70 cuts at conflicting sites were made in 29 contig sequences of the WENGAND assembly, leading to a corrected contig NGA50 of 45.91 Mb. The corrected WENGAND assembly is more contiguous than the GRCh37 reference genome (Figure 3A). The hybrid scaffolding produced 109 super-scaffold sequences with a total size of 2.83Gb and an N50 of 82Mb (Figure 4). Only 0.9% (26.29Mb) of the WENGAND sequence was not integrated into the hybrid scaffolds (short contigs). The BIONANO scaffolding of CHM13 demonstrates that the unpolished WENGAN assembly is functional and appropriate for subsequent genome analyses.

**Figure 4:**
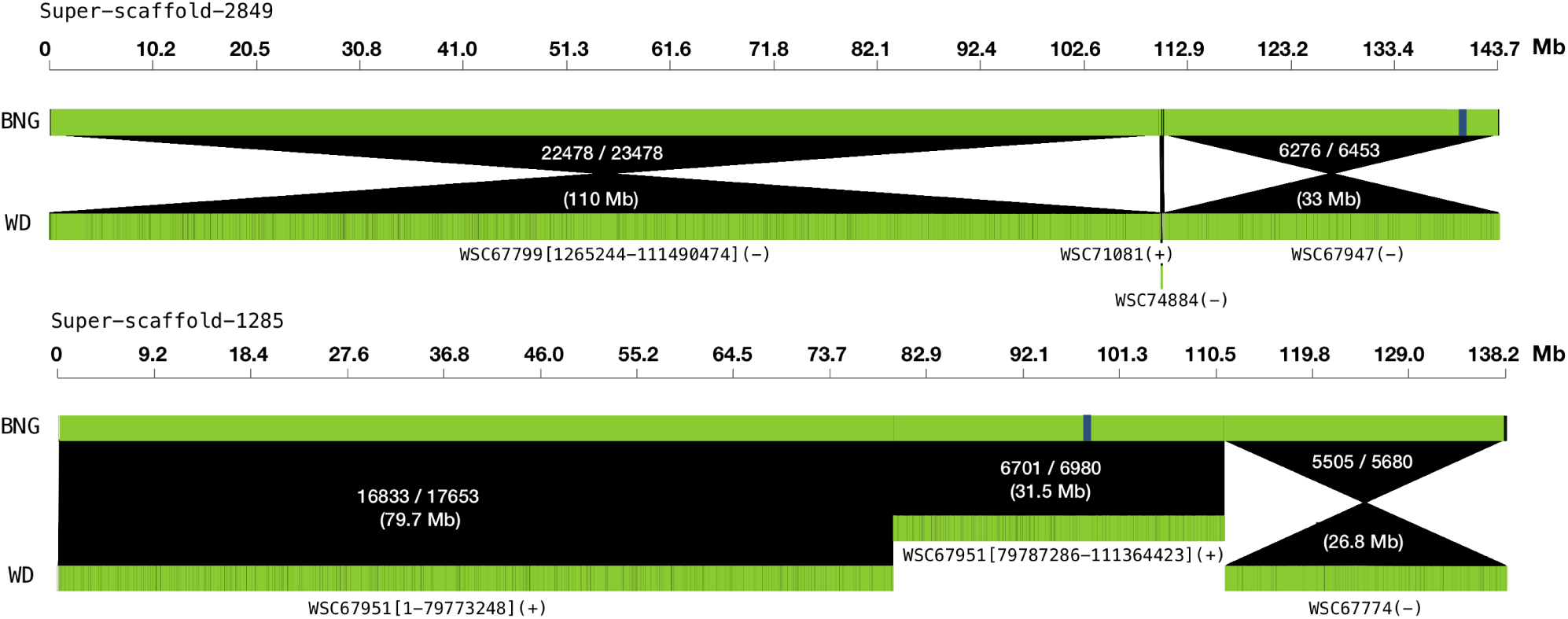
BIONANO scaffolding of the WENGAND assembly of CHM13. We show the two largest super-scaffolds produced by merging the BIONANO map (BNG) and the WENGAND contigs (WD). The name of the scaffolded WENGAND contigs is displayed (WSC). Brackets in the contig name indicate that the contig was corrected by the BIONANO map, and the numbers are the start-stop coordinates of the error-free contig region. In parenthesis, we show the contig orientation in the super-scaffold. The white text in the alignments displays the number of nicking matching sites, the total number of nicking sites in the BNG contig, and the length in megabases of the alignment. The blue bar in the BNG contigs shows examples of joins guided by the WENGAND contigs.

## 3 Discussion

In this paper, we introduced and benchmarked WENGAN, a novel and versatile genome assembler designed for enabling an efficient assembly of human-size genomes from a combination of short and long-read technologies. We demonstrated that WENGAN is the only genome assembler that optimizes the three main goals of *de novo* assembly algorithms, namely, contiguousness, consensus accuracy and computational resources. Furthermore, WENGAN is effective at shallow long-read coverage (≥15X), and in combination with ultra-long-reads generates *de novo* assemblies that surpassed the contiguity of the current human reference genome.

Additionally, we observed no significant difference in assembly quality between using the short-read platforms Illumina NovaSeq-6000 [8] or MGIseq-2000 [9] for hybrid assembly with WENGAN. Moreover, WENGAN produces high quality assemblies with any combination of short-read (NovaSeq or MGIseq-2000) and long-read (ONT MinIon/PromethION or PacBio Sequel I) technologies. In particular, we achieved a high quality human genome reconstruction for less than $1500 by hybrid assembly of ONT PromethION data (20x coverage) and BGI MGI-seq2000 (50x coverage).

Unlike current long-read assemblers, WENGAN generates functional and ready to use genome reconstructions. Polishing a large genome is not an easy task and current signal-level long-read polishers such as NANOPOLISH or ARROW need about 30k CPUs hours at 30X of genome coverage to polish a human-size genome [24, 34]. On the other hand, the consensus quality of an unpolished long-read assembly produced by FLYE or CANU is only around Q20, which corresponds to an error every 100 bases (Table S7, Table S11, Table S14, Table S15). Hence, polishing is basically mandatory for these assemblers, making their runtime a huge issue for most applications.

Nanopore-only assemblies polished with the latest algorithms [17] (Marginpolish+Helen) reach a median consensus quality of only 27.84 (or 25.4 using mean estimation, Table S14). PacBio-only assemblies with noisy continuous long reads (CLR) are slightly worse, with an estimated median QV of 27.12 (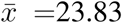, Table S15) for the diploid HG00733 sample. Thus, short-read polishing remains mandatory for Nanopore and PacBio CLR reads. Although PacBio’s HiFi-reads represent an alternative option that can mitigate the polishing step, it produces shorter long-reads 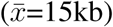, at a substantially increased price per base, and resulting in less contiguous assemblies [34]. Moreover, base calling for a 30x human genome required about 10,000 total CPU hours [34] (CCS software v4.0.0, runtime for 10 passes), making HiFi reads a problematic choice for most researchers. In summary, we found that hybrid WENGAN assemblies using Nanopore or PacBio CLR reads in combination with short-reads provide the most cost-effective and computationally efficient solution for human genome assembly to date, producing, at the same time, highly competitive assembly contiguity and quality.

We observed that the majority of the consensus errors found in WENGAN‘s assemblies are concentrated in the mate-edge sequences called from long-reads only (Figure S3). The size of such sequences ranges from 80Mb (WENGAND) to 270Mb (WENGANM) per assembly, thus reducing the amount of sequence to be polished by at least 90%. This enables local polishing of the assembled sequence and we anticipate that further polishing of the “raw” WENGAN assemblies with ad-doc algorithms may approach or exceed QV40.

WENGAN is the first genome assembler that can cope with the high-throughput of a long-read and a short-read sequencer. Although other long-read-only assemblers may have a comparable real-time execution [17](1 day), they require less accessible computational resources, more long-read coverage, and process half of the data as compared to WENGAN. Furthermore, they require computationally expensive polishing to be competitive. Still, our analyses of hard-to-assemble regions such as MHC and segmental duplications demonstrated that further algorithmic improvements are necessary for all examined assemblers. In particular, we plan to include in WENGAN novel algorithms for better exploiting pair-end information [35], trio-based phasing [36] for heterozygous genomic regions (*i.e.* MHC region), and exploiting the SSG to solve complex repeats (*i.e.* large SDs).

The anticipated rapid evolution of sequencing technologies may provide other avenues for hybrid assembly in the near future. In this paper, we enabled the effective combination of short and long-read sequencing technologies. However, the WENGAN approach also provides a natural framework to combine long-read with linked-read data and/or optical maps (BIONANO), which may lead to the assembly of ‘telomere-2-telomere’ scaffolds without the need for extra polishing and finishing methods. Therefore, we believe that WENGAN is an important step towards the democratization of *de novo* assembly of human genomes, enabling low cost - high quality genome-assembly for many research and clinical applications.

## 4 Methods

### 4.1 The WENGAN algorithm

#### 4.1.1 Short-read assembly

WENGAN can employ MINIA3 [5], ABYSS2 [6] or DISCOVARDENOVO [7] as the *de Bruijn* graph based short-read assembler. All three short-read assemblers are able to assemble a human genome in less than a day. MINIA3 and ABYSS2 were intended for low-memory assembly of large genomes. They are able to assemble human genomes using less than 40Gb of RAM [5,6]. MINIA3 is the fastest method, consuming less than 77 CPU hours to complete a human genome assembly (Supplementary Table S5). Its speed comes from the novel unipath algorithm BCALM2 [37] that uses minimizers [18] to compress quickly and with low memory the *de Bruijn* graph [37]. MINIA3 can be used iteratively to implement a multi-*k*-mer assembly approach. We used *k*-mer sizes of 41, 81 and 121 in all the WENGANM assemblies described (Supplementary Table S5). ABYSS2 uses a Bloom filter and rolling hash function as the main techniques to implement the *de Bruijn* graph-based assembly [6]. After filling the Bloom filter, ABYSS2 selects solid reads (that is, reads composed only of solid *k*-mers, namely those for which frequency(*k*) > 2) as seeds to create the unipaths. These are extended left and right by navigating in the *de Bruijn* graph until a branching vertex or a dead-end is encountered. In our benchmark tests ABYSS2 required on average 481 CPU hours to assemble a human genome (Supplementary Table S5). All the ABYSS2 assemblies were run using a Bloom Filter size of 40Gb (*B*=40G), four hash functions (*H*=4), solid *k*-mer with a minimum frequency of 3 (kc=3), *k*-mer size 96, and until the contig step only. DISCOVARDENOVO is a more specialized algorithm designed to assemble a single PCR-free paired-end Illumina library containing ≥150 base-pair reads. DISCOVARDENOVO is greedier in terms of memory than MINIA3 and ABYSS2. We observed a memory peak of 650Gb in our human assemblies (Supplementary Table S5). However, DISCOVARDENOVO better leverages the pair-end information and therefore produces the most contiguous short-read assemblies of all three tested assemblers (average contig NG50 69kb, Supplementary Table S5). All the selected short-read assemblers refine the constructed *de Bruijn* graph by removing sequencing errors and collapsing the genomic variants (SNPs,indels) to produce accurate consensus contigs [5–7].

#### 4.1.2 Pair-end pseudo-alignment as building block for genome assembly

In the same way as *k*-mers are the elemental building blocks of *de Bruijn* graph assemblers; WENGAN relies on pair-end pseudo-aligments as the elemental building blocks for the *de novo* assembly. We recently introduced an alignment-free method called FAST-SG [27] that uses unique *k*-mers to compute a pseudo-alignment of pair-end reads from long or short-reads technologies. Here, we present its successor, which we called FASTMIN-SG, which implements the same ideas of FAST-SG but using minimizers [18] and chaining with the MINIMAP2 API [38]. The uniqueness of the pseudo-alignment is now determined using the MINIMAP2 mapping quality score which gives a higher score to a primary chain when its best secondary chain has a weak pseudo-alignment.

To perform a pseudo-alignment of pair-ends from short-read sequencing technologies, we use (10,21)-minimizers for querying and indexing. We discard pair-end pseudo-alignments when one of the mates has a mapping quality score ≤ 30 or covers ≤ 50% of the read bases. For mapping synthetic pair-ends extracted from long-read technologies, we use (5,20)-minimizers and a read length of 250 base pairs. A synthetic pair-end is a fragment of length *d* for which we have access to the long-read of origin, the position of the fragment in the long-read and the inner long-read sequence between both mates of the synthetic fragment. All the synthetic fragments are extracted from the long-reads using a moving window of 150bp in forward-reverse orientation. We create a spectrum of synthetic mate-pair libraries (Figure S2) by extracting pair-ends at different distances. The range of distances depends on the long-read lengths but go from 0.5kb to a maximum of 500kb with ultra-long nanopore reads. For noisy PacBio reads, we use homopolymer compressed *k*-mers [38] for indexing and querying the synthetic pair-ends. We discard synthetic pair-end alignments when one of the mates has a mapping quality score ≤ 40 or covers ≤ 65% of the synthetic read bases. The information associated to the long-read of each synthetic pair is stored in the read names for computing approximate long-read alignments later. FASTMIN-SG, like MINIMAP2, uses presets to modify multiple parameters, thus simplifying its usability. Currently, it has presets for raw PacBio reads (pacraw), raw (ontraw) and ultra-long (ontlon) Oxford Nanopore reads, and pair-ends (shortr) from short-read technologies (supporting Illumina or BGI). The pseudo-alignments are reported in SAM format.

#### 4.1.3 Detection and split of chimeric short-read contigs

The *de Bruijn* graph is complex around repeat sequences, and short-read assemblers can choose wrong paths while traversing such complex regions, thus leading to chimeric contigs (Figure S1). To detect potential chimeric contigs not supported by the short-reads, we map the pair-end reads back to the assembled short-read contigs using FASTMIN-SG (preset shortr). From the pair-end pseudo-alignments, we infer the average 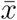 and standard deviation *σ* of the insert-sizes distribution of the genomic library. Then, pair-ends mapped within contigs at the expected orientation and distance 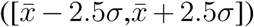 are transformed into physical fragments. For each contig, we create an array of length equal to the contig length, and the contig fragments are used to increase the physical coverage of the contig bases. We then scan the physical coverage array base by base to detect Low Quality Intervals (LQI) that have fragment coverage below a minimum depth threshold (def:7). LQIs are classified according to their contig location as internal, start, end or whole. Finally, contigs are trimmed/split at the boundaries of the LQIs.

#### 4.1.4 The Synthetic Scaffolding Graph (SSG)

We build on top of the work of Huson *et al.* [28] to extend the scaffolding graph formulation and create the Synthetic Scaffolding Graph (SSG). In brief, the contig scaffolding problem was defined by Huson *et al.* [28] as the determination of an order and orientation of a set of contigs that maximizes the amount of satisfied mate-pair links. The scaffolding graph *G* = (*V, E*), with vertex set *V* and edge set *E*, is a weighted, undirected multi-graph, without self-loops [28]. Each contig *C*_*i*_ is modeled by two vertices (*v, w*) and an undirected contig-edge (*e*). The length of *e* is set to the contig length *l*(*C*_*i*_). The contig orientation is represented by associating each of the contig ends to one of the two vertices (*i.e. tail*(*C*_*i*_) = *v* and *head*(*C*_*i*_) = *w*). Then traversing from *tail*(*C*_*i*_) → *head*(*C*_*i*_) or *head*(*C*_*i*_) → *tail*(*C*_*i*_) implies forward or reverse contig orientation, respectively. Now, consider a pair of mate-reads *f* and *r* originated from a synthetic mate-pair library with mean insert size 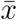, standard deviation *σ* and orientation forward-reverse that uniquely matches two different contigs *C*_*i*_ and *C*_*j*_. The uniquely mapped mate-pair induces a relative orientation and approximate distance between the two contigs. Such information is represented by adding a mate-edge *e* into the graph. The length of the mate-edge *e* is computed by subtracting from the expected mate-pair distance 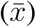 the amount of overlap that each contig has with the mate-pair considering the read mapping orientations: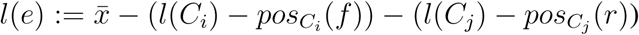. Moreover, the standard deviation *σ*(*e*) of each mate-edge *e* is set equal to the standard deviation of the synthetic mate-pair library. If there is more than one mate-edge *e* between the same ends of two contigs *C*_*i*_ and *C*_*j*_, we can bundle [28] the mate-edge *e* by computing from the set of mate-edges *e*_1_, *e*_2_, …, *e*_*n*_, the length of *e* as *l*(*e*): = *p/q* and its deviation as 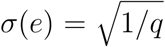 where: 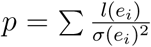 and 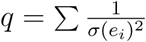 [28]. Additionally, the weight *w*(*e*) of a bundled mate-edge *e* is set to 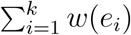, otherwise to 1.

The **Synthetic Scaffolding Graph (SSG)** is an edge-bundled scaffolding graph *G* = (*V, E*), built from a **spectrum** of synthetic mate-pair libraries, where there is an edge-labelling function (*F*) that maps the long reads to the edges through the synthetic mate-pair pseudo-alignments.

#### 4.1.5 Computing approximate long-read overlaps with the SSG

Consider a mate-edge *e* from *v* to *w* that is connected by an alternative path *P* = (*m*_1_, *c*_1_, *m*_2_, …, *m*_*k*_) of mate-edges (*m*_1_, *m*_2_, …), contig-edges (*c*_1_, *c*_2_, …) and long-read labels *F* (*P*) = (*F* (*m*_1_), *F* (*c*_1_), *F* (*m*_2_), …, *F* (*m*_*k*_)). We can compute the path length *l*(*P*) and its standard deviation *σ*(*P*) as follows [28]: *l*(*P*): = Σ *l*(*m*_*i*_)+ Σ *l*(*c*_*i*_) and 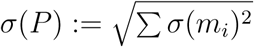. A mate-edge *e* from *v* to *w* can be transitively reduced on the path *P* if *e* and *P* have similar lengths and the long-read labels of *e* are coherent with every edge *e*_*i*_ of *P*: |*l*(*e*) −*l*(*P*)| ≤ 4 max(*σ*(*e*), *σ*(*P*)) and *F* (*e*) ⊂ *F* (*e*_*i*_) ∀ *e*_*i*_ ∈ *P*. If this is the case, then the alternative path *P* is long-read coherent with the mate-edge *e* and represents an approximate overlap of length *l*(*P*) among all the long-reads composing the mate-edge *e* (*F* (*e*)). We store the overlap information by removing *e* from the SSG and incrementing the weight of every mate-edge *m*_*i*_ in *P* by *w*(*e*). Before starting the computation of approximate long-read overlaps, the mate-edges are sorted by ascending length *l*(*e*) and the set of biconnected components of the SSG is computed. Alternative long-read-coherent path search takes place inside each biconnected component. In practice, we use a depth-first search algorithm to enumerate all the long-read-coherent paths of a given mate-edge *e*. At each edge extension, we extend the path only if the new added edge is long-read-coherent with the given mate-edge *e* (*F* (*e*) ⊂ *F* (*e*_*i*_)). Contig-edges associated to repeat sequences are masked during path extension. We stop searching when the size of a partial path *P* is larger than 80 vertices or its length is longer than expected (*l*(*P*) > *l*(*e*) and |*l*(*e*) −*l*(*P*)| > 4 max(*σ*(*e*), *σ*(*P*))). If there is more than one long-read-coherent path, we choose the path having the maximum number of hits from the long-reads supporting the given mate-edge *e*. For very long mate-edges (*l*(*e*) ≥ 100*kb*), we stop searching if we find more than 100 long-read-coherent paths. All the selected long-read-coherent paths are stored in a path database for later use.

The final SSG graph is created by performing first bundling and then transitive reduction (approximate long-read overlaps) of mate-edges. From now on, we will refer to this simply as the *reduced SSG*.

#### 4.1.6 Generation of the assembly backbone with the SSG

Computation of approximate long-read overlaps allows to solve the scaffolding problem using all the synthetic mate-pair libraries simultaneously. Given the reduced SSG, our goal is to determine an optimal set of vertex disjoint paths covering all the contig-edges with a maximum total weight of the mate-edges. Since this optimization problem is NP-hard, we use Edmond’s maximum weighted matching approximation algorithm that guarantees to find an optimal solution with a worst-case performance ratio *r* = *W* (*S*)*/W* (*G*) > 2*/*3 [29]. The matching algorithm implementation is based on an extensive use of priority queues, leading to an *O*(*V E* log(*V*)) time complexity [39,40]. All the contig-edges, as well as the mate-edges associated to repetitive contigs or having a weight smaller than 5, are masked during the matching cover step. After computing the matching cover, all the contig-edges are added to the matching cover solution and we use a depth-first-search approach to detect simple cycles. If such cycles are found, the set of biconnected components of the graph is computed and simple cycles are destroyed by removing the mate-edge of lowest weight in each biconnected component. In practice, the matching cover solutions contain few cycles (< 10 on human genomes) and we observed performance-ratios higher than *r* > 0.8. The set of optimal simple paths (scaffolds) is what we call the *assembly backbone*.

#### 4.1.7 Validation of the assembly backbone with the SSG

We validate the assembly backbone using the physical genomic coverage obtained from the computation of the approximate long-read overlaps. The key idea is to identify suspicious mate-edges *e* (corresponding to potentially incorrect joins) not supported/covered by long-read overlaps longer than *O* (by default *O ≥* 20*kb*). We first assign genomic coordinates to each line *L*_*i*_ = (*c*_1_, *m*_2_, …, *m*_*k*−1_, *c*_*k*_) from 1 to *l*(*L*_*i*_), taking into account the orientation of the contig-edges and the ordering and distance provided by the matched mate-edges. In a second step, all the contig-edges are converted into physical genomic fragments as well as the mate-edges spanned by long-reads longer than *O*. In a third step, if the vertices *v, w* of a reduced mate-edge *e* belong to the same line *L*_*i*_ and the length of *l*(*e*) is longer than *O*, we create a new simple path *pf* that goes from *v* to *w* in the line *L*_*i*_. The new simple path *pf* is converted into a physical genomic fragment *f* only if the length of *pf* is similar to the length of the reduced mate-edge *e*, that is, if |*l*(*e*) *-l*(*pf*)| ≤ 4 max(*σ*(*e*), *σ*(*pf*)). If that is the case, the simple path *pf* increases the physical genomic coverage of the line *L*_*i*_. In a forth step, once the physical genomic coverage of all the lines *L*_*i*_ have been computed, we look for all the intervals inside a scaffold having a lack of physical coverage at the mate-edge locations and we tag such mate-edges as potentially erroneous joins. A line *L*_*i*_ is split at potential error joins only if the number of long-reads supporting the suspicious mate-edge *e* is less than *mlr* (default *mlr ≤* 4). In practice, we observe that the physical path coverage of human assemblies is around 20X (30X long-read coverage), thus usually less than 200 mate-edges are removed.

#### 4.1.8 Gap filling with the SSG

A property of the SSG is that all the mate-edges are spanned by at least one long-read. Therefore, after construction and validation of the assembly backbone, we proceed to create a consensus sequence for each of the matched mate-edges. We start by ordering the lines by decreasing length, which imposes a global order to the mate-edges and consequently to the long-read sequences. For each mate-edge *e*, we select the *N* best long-reads (default: 20) spanning *e*. The long-read selection is done by counting, with the edge-labeling function *F* (*e*), the number of synthetic mate-pairs contributed by the long-read *l*_*i*_ to compose the mate-edge *e*. This means that, the more synthetic mate-pairs are contributed by long-read *l*_*i*_, the greater is the confidence that the long-read *li* spans *e*. All the selected long-read sequences are sorted according to the mate-edges order using an external merge sort algorithm to create a long-read sequence database. Following the long-read database creation, we build a consensus sequence for each mate-edge using the partial order alignment graph [30]. For each mate-edge, we select the long-read contributing the most synthetic mate-pairs as consensus template, then the remaining long-reads spanning *e* are aligned to the template using a fast implementation of Myers’ bit-vector algorithm [31]. The long-read alignments are scanned to partition the long-reads into non-overlapping windows of size *w* (by default 500bp) on the template sequence. The long-read chunks that have an average identity lower than 65% are removed from the corresponding windows. The purpose is to use high-quality alignments to build the template consensus. For each window *w*, we call the consensus sequence using a SIMD accelerated implementation of the partial order alignment graph [15]. The mate-edge consensus is built by joining the window sequences. Finally, the corresponding contig-ends are aligned (using once again Myers’ bit-vector algorithm) to the mate-edge consensus sequence to determine the correct mate-edge sequence boundaries, thus filling the gap between the two contig-edges.

#### 4.1.9 Polishing with the SSG

Since not all the contig-edges are part of the assembly backbone (as is the case for the contig-edges related to repeats or short sequences), we can use them to improve the consensus base accuracy of the mate-edge sequences. To this end, we use two polishing strategies, one based on the SSG and a second based on pairwise alignments. The graph polisher uses the reduced SSG to find alternative long-read-coherent paths as before, but masking the contig-edges composing the assembly backbone. Since now we navigate on more tangled parts of the SSG (unmasked repeat sequences), we limit the path search to a maximum of 5 million iterations on each mate-edge. Once a long-read-coherent path has been found, we align the contig-edges (with the proper orientation) to the mate-edge sequence using Myers’ bit-vector algorithm [31]. Then, the alignments are trimmed as a function of the average long-read-depth of the mate-edge consensus sequence. We thus expect a minimum identity between 80% and 99% when the average long-read-depth of the consensus sequence is between 1 and 20, respectively. If a contig-edge maps with an identity higher than the expected and the alignment covers at least 75% of the contig-edge, we replace the corresponding mate-edge aligned sequence by the contig-edge aligned sequence, thus polishing the mate-edge sequence. The alignment polisher searches for matches between the singleton contig-edges and all the mate-edge consensus sequences. In brief, we first index all the mate-edge consensus sequences using (5,17)-minimizers [18]. Minimizers are stored in a hash table and the ones having a frequency higher than 1000 are excluded. The (5,17)-minimizers of the contig-edges are scanned on the mate-edge sequence index to collect HSPs or exact (5,17)-minimizer matches. HSPs are sorted by mate-edges and hits are identified by finding the longest strictly increasing subsequence (co-linear chain) between the contig and the mate edges. After collecting all the hits, we use a greedy algorithm to determine a layout of contig-edge hits along the mate-edge sequence. The greedy algorithm starts by sorting the contig-edge hits by number of minimizer matches and then adds the hits to the layout only if there is no overlap with a previously added hit. We then proceed as in the graph polisher to align and polish the mate-edge sequence using the best-hit layout. Finally, WENGAN outputs the sequence of each line plus the sequence of contig-edges (>5kb) not used in the polishing steps.

### 4.2 Assembly validation

Genome assemblies generated by WENGAN and other assemblers were assessed by whole genome alignment to the human reference genome using the MINIMAP2 [38] program and a procedure similar to the one described recently by Ruan *et al.* [14]. To determine breakpoints, a maximum of 5% divergence in the alignments was allowed (MINIMAP2 options: –paf-no-hit - cxasm20 -r2k -z1000,500) and then alignments longer than 50kb were scanned by PAFTOOLS (option asmstat -q50000 -d.1) to collect several assembly metrics (*i.e.* NG50, NGA50, largest alignment block and others). The consensus quality was determined by computing a more stringent alignment allowing a maximum of 1% divergence (MINIMAP2 options: cxasm10 –cs -r2k), then alignments longer than 50kb were scanned by PAFTOOLS (option call -l50000 -L50000) to call Single Nucleotide Variants, insertions and deletions. Additionally, we used the 100-mers completeness analysis to assess with an alignment-free method the consensus quality of the genome assemblies using the KMC [41] *k*-mer counter (Version 3.1.0). The GRCh37 (hs37d5) reference genome has a total of 2,825,844,566 distinct 100-mers and those were intersected with the 100-mers of the genome assemblies using the KMC_TOOLS utility (option intersect -ci1 -cx1000). A QUAST [32] (Version: 5.0.2) analysis was run with the options min-identity 80 and fragmented to validate the NA12878 assemblies (rel5) using hs37d5 as reference and the HG0075, NA243875 and CHM13 assemblies using the GRCh38 reference (autosomes plus X and Y). The gene completeness of the genome assemblies was assessed with the BUSCO [42] program (Version 3.0.2) using the MAMMALIA ODB9 gene set (4104 BUSCO groups). The single plus duplicated complete BUSCO gene counts are reported. The consensus quality of the genome assemblies was determined by aligning orthogonal BAC or Fosmid sequences data (Table S4). The statistics were computed considering fully resolved BAC/Fosmid only. The BAC/Fosmid consensus quality analysis was performed using the BACVALIDATION tool (https://github.com/skoren/bacValidation). The amount of Segmental Duplication (SD) resolved by the genome assemblies of CHM13, HG00733 and NA24385 was determined using SEGDUPPLOTS [33] (https://github.com/mvollger/segDupPlots). SEGDUPPLOTS aligns the assembled contigs to GRCh38 and considers an SD as resolved when the aligned contig extends the SD flanking sequences by at least 50kb. Finally, the WENGAND assembly of CHM13 was validated and scaffolded using the hybridScaffold.pl program (BIONANOSolve3.4 06042019a) (with the options -c hybridScaffold DLE1 config.xml -B 2 -N 2) and the BIONANO map assembled by the T2T-consortium [25].

### 4.3 Hybrid polishing of FLYE assemblies

We polished the FLYE assemblies of NA12878 using the same sequencing reads employed in the WENGAN assemblies. We used two rounds of long-read polishing with RACON [15] followed by three rounds of short-read polishing with NTEDIT [22]. The commands executed as well as the consensus quality improvement after each round of polishing are provided in the Supplementary Material (Table S8 and Table S12).

### 4.4 Genome sequencing of NA12878

The genomic DNA from the GM12878 human cell line was purchased from the Coriell Institute (cat. no. NA12878).

#### 4.4.1 MGI sequencing

Library preparation for the NA12878 sample was performed with the MGIEasy DNA Library Prep Kit V1.1 (MGI, 940-200022-00) following the manufacturer’s instructions. Briefly, 1*µ*g of genomic DNA at a concentration of 12.5ng/*µ*L was fragmented with an E220 Covaris program optimized to yield fragments of 450bp average length. A double-sized selection was performed with AMPure XP beads (Beckman Coulter) at 0.52X ratio followed by a 0.15X ratio as recommended by MGI. A total of 50ng of fragmented DNA was used for the end repair and A-tailing reaction following the manufacturer’s instructions. A set of adapters with 8 barcodes were ligated to the repaired DNA for one hour at 23*°*C. After purification with AMPure XP beads (Beckman Coulter) at a 0.5X ratio, the DNA was subjected to PCR enrichment following the manufacturer’s instructions. A total of 330ng of PCR product was hybridized with the Split Oligo (MGI, 940-200022-00) for the circularization step followed by digestion. Circularized ssDNA was purified with Library Purification Beads (MGI, 940-200022-00) and quantified with an ssDNA assay on a Qubit 3 fluorometer (Thermo Fisher). For the linear amplification to generate DNA nanoballs (DNBs), 75fmol of circularized ssDNA were used. The DNB library was loaded in a single lane and sequenced on a MGISEQ-2000 instrument with a paired-end modus and read length of 150bp with the MGISEQ-2000RS High-throughput Sequencing Set PE150 (MGI, 1000003981) according to (DNBs) manufacturer’s instructions.

#### 4.4.2 Illumina sequencing

Library was prepared using the TruSeq DNA PCR-Free Library Prep kit (Illumina, FC-121-3001) following the TruSeq DNA PCR-free reference guide (Illumina, 1000000039279v00). Briefly, 1*µ*g of genomic DNA was used for fragmentation on an E220 Covaris to yield insert sizes of 350bp. The DNA was end-repaired, adenylated and subjected to adapter ligation as described in the reference guide. The library was quantified using the KAPA Lib Quantification Kit (Roche, LB3111) and dsDNA HS assay (Qubit). The average fragment size was estimated with a HS DNA kit (Agilent) on a 2100 Bioanalyzer (Agilent). A S2 flow cell was loaded with 2.2nM on a NovaSeq6000 instrument to generate 2×151 paired-end reads.

#### 4.4.3 Oxford Nanopore sequencing

Three flow cells were run with the sample NA12878. One flow cell was loaded with a library prepared from unsheared genomic DNA. For the additional two sequencing runs, [14*µ*g of] NA12878 genomic DNA was mechanically sheared with Megaruptor 3 (Diagenode) [at a concentration of 70 ng/*µ*L in a volume of 200*µ*L] with manufacturer’s recommended speed to get sheared DNA with an average fragment length of 30Kb. Size selection was performed with Blue Pippin*™* (Sage Science) to remove fragments shorter than 10Kb using a 0.75% agarose cassette, the S1 marker and a high-pass protocol (Biozym, 342BLF7510). A further clean-up with AMPure XP beads (Beckman Coulter) on the size-selected DNA was performed at a 1X ratio for one library. Fragment size was assessed with the gDNA 165kb Analysis Kit on a FemtoPulse (Agilent) and the concentration of DNA was assessed using the dsDNA HS assay on a Qubit 3 fluorometer (Thermo Fisher). For each of the three sequencing runs, one library was prepared with the SQK-LSK109 Ligation Sequencing kit (ONT) per flow cell following the instructions of the 1D Genomic DNA by Ligation protocol from Oxford Nanopore Technologies. Briefly, 1.1 to 1.3*µ*g of genomic DNA was used for the DNA repair reaction with NEBNext Ultra II End Repair/dA-Tailing Module (New England Biolabs, E7546S) and NEBNext FFPE DNA Repair Module (NEB, M6630S). Upon clean-up with AMPure XP beads (Beckman Coulter) at 1X ratio, the end-repaired DNA was incubated for one hour at room temperature with Adapter Mix (ONT, SQK-LSK109), Ligation Buffer (ONT, SQK-LSK109) and NEBNext Quick Ligation Module (NEB, E6056S). The ligation reaction was purified with AMPure XP beads (Beckman Coulter) at a 0.4X ratio and L Fragment Buffer (ONT, SQK-LSK109). A total of 600ng (25fmol) of the generated libraries were loaded into the flow cell (FLO-PR002) on a PromethION instrument (ONT) following the manufacturer’s instructions.

### 4.5 Availability

The WENGAN software and all the WENGAN assemblies described in the present manuscript are freely available at https://github.com/adigenova/wengan. The NovaSeq6000, MGISEQ-2000RS and PromethION sequence data of NA12878 was submitted to the European Nucleotide Archive (ENA) under the BioProject XXXX.

## Acknowledgements

The authors are grateful to Vincent Lacroix for helpful comments on initial versions of the manuscript. We would like to thank the NCCT members Antje Schulze Selting and Christina Engensser for excellent technical assistance.

## Author contributions

ADG devised the original ideas for WENGAN, developed, implemented and bench-marked WENGAN. MFS guided the development of WENGAN. EBA performed sequencing of NA12878 and primary quality control of sequenced data. ADG wrote the initial version of the manuscript. MFS and SO improved initial versions of the manuscript. All authors were involved in discussions and revisions of the project and manuscript.

