## Supplementary material for "WENGAN: Efficient and high quality hybrid *de novo* assembly of human genomes"

### 1 Genomes assemblies

#### 1.1 Short-read assemblies

##### 1.1.1 ABYSS

```
#Abyss version 2.1.5

#NA12878 2x150bp HiSeq 2500
abyss-pe name=NA12878-abyss-ILL60X150 np=20 k=96 lib="pea peb" pea="BH88WKADXX.R1.fastq.gz BH88WKADXX.R2.fastq.gz"
      peb="AH81VLADXX.R1.fastq.gz AH81VLADXX.R2.fastq.gz" B=40G H=4 kc=3 v=-v contigs
#NA12878 2x150bp NovaSeq
abyss-pe name=NA12878-abyss-NS np=20 k=96 lib="pea" pea="S22_L001_R1_001.fastq.gz S22_L001_R2_001.fastq.gz" B=40G H=4 kc=3 v=-v
      contigs
# NA12878 2x150bp MGI-2000
abyss-pe name=NA12878-abyss-MGI np=20 k=96 lib="pe1 pe2" pe1="NA12878EBA.bgi.fwd.fastq.gz NA12878EBA.bgi.rev.fastq.gz"
      pe2="MGISEQ.sample.fwd.gz MGISEQ.sample.rev.gz" B=40G H=4 kc=3 v=-v contigs
```

##### 1.1.2 DISCOVARDENOVO

```
#Discovar version discovarexp-51885
DISCO=/path/DiscovarExp
export MALLOC_PER_THREAD=1
#NA12878 2x250bp HiSeq 2500
${DISCO} READS="SRR891258_{1,2}.fastq.gz,SRR891259_{1,2}.fastq.gz" NUM_THREADS=44 OUT_DIR=60XNA12878
#NA12878 2x150bp NovaSeq
 ${DISCO} READS="S22_L001_R{1,2}_001.fastq.gz" NUM_THREADS=44 OUT_DIR=NA12878NS
# NA12878 2x150bp MGI-2000
${DISCO} READS="NA12878EBA.bgi.R{1,2}.fastq.gz,MGISEQ.sample.R{1,2}.fastq.gz" NUM_THREADS=44 OUT_DIR=NA12878MGI
# NA24385
# prior to assembly the reads shorter than 100 bp were discarded with fastp
fastp -l 100 -i D1_S1S2_R1.fastq.gz -I D1_S1S2_R2.fastq.gz -o D1_S1S2T_R1.fastq.gz -O D1_S1S2T_R2.fastq.gz
${DISCO} READS="D1_S1S2T_R{1,2}.fastq.gz" NUM_THREADS=44 OUT_DIR=60XNA24385
# HG00733
${DISCO} READS="SRR5534476_{1,2}.fastq.gz,SRR5534475_{1,2}.fastq.gz" NUM_THREADS=44 OUT_DIR=60XHG00733
#CHM13
${DISCO} READS="SRR3189741_{1,2}.fastq.gz,SRR3189742_{1,2}.fastq.gz" NUM_THREADS=44 OUT_DIR=60XCHM13
```

##### 1.1.3 MINIA3

```
# Minia 3, git commit 017d23e
#NA12878 2x150bp HiSeq 2500
echo BH88WKADXX.R1.fastq.gz BH88WKADXX.R2.fastq.gz AH81VLADXX.R1.fastq.gz AH81VLADXX.R2.fastq.gz | xargs -n 1 > reads.txt
# the script run-minia.pl runs a mult-K assembly with kmer-sizes 41,81,121 and min k-mer frequencies of 2,2,2 respectively.
perl run-minia.pl -a reads.txt -c 20 -p NA12878-Hiseq
#NA12878 2x150bp NovaSeq
```

```
echo S22_L001_R1_001.fastq.gz S22_L001_R2_001.fastq.gz | xargs -n 1 > reads.txt
perl run-minia.pl -a reads.txt -c 20 -p NA12878-NS
# NA12878 2x150bp MGI-2000
echo NA12878EBA.bgi.fwd.fastq.gz NA12878EBA.bgi.rev.fastq.gz MGISEQ.sample.fwd.gz MGISEQ.sample.rev.gz | xargs -n 1 > reads.txt
perl run-minia.pl -a reads.txt -c 20 -p NA12878-MGI
```

The script *run-minia.pl* is part of the WENGAN code (directory aux_scripts/run-minia.pl).

#### 1.2 WENGAN assemblies

##### 1.2.1 WENGAN assemblies of NA12878, NA24385, HG00733, and CHM13

```
#NA12878
#WenganA
wengan.pl -x ontlon -a A -s AH81VLADXX.R1.fastq.gz,AH81VLADXX.R2.fastq.gz,BH88WKADXX.R1.fastq.gz,BH88WKADXX.R2.fastq.gz -l
    na12878.rel5.fastq.gz -p na12878Wa -t 20 -g 3000
#WenganD high memory machine
wengan.pl -x ontlon -M 5000 -a D -s SRR891259_1.fastq.gz,SRR891259_2.fastq.gz,SRR891258_1.fastq.gz,SRR891258_2.fastq.gz -l
    na12878.rel5.fastq.gz -p na12878Wd -t 44 -g 3000
#WenganM
perl wengan.pl -x ontlon -a M -s AH81VLADXX.R1.fastq.gz,AH81VLADXX.R2.fastq.gz,BH88WKADXX.R1.fastq.gz,BH88WKADXX.R2.fastq.gz -l
    na12878.rel5.fastq.gz -p na12878Wm -t 20 -g 3000

#NA24385
#upto 500kb
LIBS=500,1000,2000,3000,4000,5000,6000,7000,8000,10000,15000,20000,30000,40000,50000,60000,70000,80000,90000,100000,120000,150000,
    180000,200000,250000,300000,350000,400000,450000,500000
#run the WenganD pipeline from the given Discovar contigs
perl wengan.pl -x ontlon -M 5000 -P 100000 -a D -s D1_S1S2T_R1.fastq.gz,D1_S1S2T_R2.fastq.gz -l ultra-long-ont.fastq.gz -p
    NA24385Wd -t 20 -g 3000 -c 60XNA24385.disco.fa -i ${LIBS}

#HG00733
#upto 80Kb
LIBS=500,1000,2000,3000,4000,5000,6000,7000,8000,10000,15000,20000,30000,40000,50000,60000,70000,80000
#run the WenganD pipeline from the given Discovar contigs
perl wengan.pl -x pacraw -a D -s SRR5534475_1.fastq.gz,SRR5534475_2.fastq.gz,SRR5534476_1.fastq.gz,SRR5534476_2.fastq.gz -l
    SRR7615963_subreads.fastq.gz -p HG00733Wd -t 20 -g 3000 -c HG00733.ILL250.DISCOVAR.fa -i ${LIBS}

#CHM13
#upto 500kb
LIBS=500,1000,2000,3000,4000,5000,6000,7000,8000,10000,15000,20000,30000,40000,50000,60000,70000,80000,90000,100000,120000,150000,
    180000,200000,250000,300000,350000,400000,450000,500000
#run the WenganD pipeline from the given Discovar contigs
perl wengan.pl -x ontlon -M 4000 -P 100000 -a D -s
    SRR3189741_1.fastq.gz,SRR3189741_2.fastq.gz,SRR3189742_1.fastq.gz,SRR3189742_2.fastq.gz -l ont.rel2.fastq.gz -p CHM13Wd -t
    20 -g 3000 -c 60XCHM13L.disco.fa -i ${LIBS}
```

##### 1.2.2 WENGAN assemblies of NA12878 at different long-read coverage

The command used for WENGAN assemblies with MGI+ONT data are shown. Identical commands were used for WENGAN assemblies from ILL+ONT data.

```
# WenganA
#10X
perl wengan.pl -M 1000 -N 2 -x ontraw -a A -s short.reads -l 10X.fastq.gz -p 10X -g 3000 -t 20 2>10X.err > 10X.log
#15X
perl wengan.pl -M 1000 -N 3 -x ontraw -a A -s short.reads -l 15X.fastq.gz -p 15X -g 3000 -t 20 2>15X.err > 15X.log
#20X
perl wengan.pl -M 1000 -N 4 -x ontraw -a A -s short.reads -l 20X.fastq.gz -p 20X -g 3000 -t 20 2>20X.err > 20X.log
#25X
perl wengan.pl -M 1000 -N 5 -x ontraw -a A -s short.reads -l 25X.fastq.gz -p 25X -g 3000 -t 20 2>25X.err > 25X.log
#30X
perl wengan.pl -M 1000 -N 5 -x ontraw -a A -s short.reads -l 30X.fastq.gz -p 30X -g 3000 -t 20 2>30X.err > 30X.log

# WenganD
#10X
perl wengan.pl -N 2 -x ontraw -a D -s short.reads -l 10X.fastq.gz -p 10X -g 3000 -t 44 2>10X.err > 10X.log
#15X
perl wengan.pl -N 3 -x ontraw -a D -s short.reads -l 15X.fastq.gz -p 15X -g 3000 -t 44 2>15X.err > 15X.log
#20X
perl wengan.pl -N 4 -x ontraw -a D -s short.reads -l 20X.fastq.gz -p 20X -g 3000 -t 44 2>20X.err > 20X.log
#25X
perl wengan.pl -N 5 -x ontraw -a D -s short.reads -l 25X.fastq.gz -p 25X -g 3000 -t 44 2>25X.err > 25X.log
#30X
perl wengan.pl -N 5 -x ontraw -a D -s short.reads -l 30X.fastq.gz -p 30X -g 3000 -t 44 2>30X.err > 30X.log

# WenganM
#10X
perl wengan.pl -M 1000 -N 2 -x ontraw -a M -s short.reads -l 10X.fastq.gz -p 10X -g 3000 -t 20 2>10X.err > 10X.log
#15X
perl wengan.pl -M 1000 -N 3 -x ontraw -a M -s short.reads -l 15X.fastq.gz -p 15X -g 3000 -t 20 2>15X.err > 15X.log
#20X
perl wengan.pl -M 1000 -N 4 -x ontraw -a M -s short.reads -l 20X.fastq.gz -p 20X -g 3000 -t 20 2>20X.err > 20X.log
#25X
perl wengan.pl -M 1000 -N 5 -x ontraw -a M -s short.reads -l 25X.fastq.gz -p 25X -g 3000 -t 20 2>25X.err > 25X.log
#30X
perl wengan.pl -M 1000 -N 5 -x ontraw -a M -s short.reads -l 30X.fastq.gz -p 30X -g 3000 -t 20 2>30X.err > 30X.log
```

##### 1.3 Flye assemblies

###### 1.3.1 Flye assemblies at different long-read coverage

```
# Flye v2.5
#10X
flye --nano-raw 10X.fastq.gz -o 10X.flye -t 44 -g 3g
#15X
flye --nano-raw 15X.fastq.gz -o 15X.flye -t 44 -g 3g
#20X
flye --nano-raw 20X.fastq.gz -o 20X.flye -t 44 -g 3g
#25X
flye --nano-raw 25X.fastq.gz -o 25X.flye -t 44 -g 3g
#30X
flye --nano-raw 30X.fastq.gz -o 30X.flye -t 44 -g 3g
```

#### 2 Polishing Flye assemblies

##### 2.1 Polishing with short and long reads

Flye assemblies of NA12878 were polished using Racon and ntEdit. In particular, two rounds of long-read polishing with Racon were performed, followed by three rounds of short-read polishing with ntEdit. The commands executed were the following:

```
#Polishing of the Flye assembly with 40X nanopore reads (rel5) and 50X of short illumina reads.
make -f polish.mk PREF=na12878.flye.racon ASM=FLYE.NA12878.fa CPU=44 READS=na12878.rel5.fastq.gz SREADS="AH81VLADXX.R1.fastq.gz
     AH81VLADXX.R2.fastq.gz BH88WKADXX.R1.fastq.gz BH88WKADXX.R2.fastq.gz" all
#Polishing of the Flye assembly at 30X coverage with flip-flop called nanopore reads and Illumina Novaseq short-reads
make -f polish.mk PREF=na12878.flye30x.racon ASM=na12878.flye30X.fa CPU=44 READS=30X.fastq.gz SREADS="S22_L001_R1_001.fastq.gz
     S22_L001_R2_001.fastq.gz" all
```

The makefile *polish.mk* contains the following instructions:

```
.DELETE_ON_ERROR:
#racon version v1.4.9
RACON=/path/racon
#minimap2 version 2.15-r905
MM=/path/minimap2
# nthits version 0.1.0
NTH=/path/nthits
# ntedit version 1.2.3
NTE=/path/ntedit
TIME=/path/time
#LONG-READ POLISHING
#first round of long-read polishing
$(PREF).r1.paf:
	${TIME} -v -o $(PREF).r1.mm.time ${MM} -x map-ont -t ${CPU} ${ASM} ${READS} > $@
$(PREF).r1.fa:$(PREF).r1.paf
	${TIME} -v -o $(PREF).r1.racon.time ${RACON} -u -t ${CPU} ${READS} $< ${ASM} > $@ 2> $(PREF).r1.racon.log
#second round of long-read polishing
$(PREF).r2.paf:$(PREF).r1.fa
	${TIME} -v -o $(PREF).r2.mm.time ${MM} -x map-ont -t ${CPU} $< ${READS} > $@
$(PREF).r2.fa:$(PREF).r2.paf
	${TIME} -v -o $(PREF).r2.racon.time ${RACON} -t ${CPU} ${READS} $< $(PREF).r1.fa > $@ 2> $(PREF).r2.racon.log
#SHORT-READ POLISHING
$(PREF).nthits_k60.bf:$(PREF).r2.fa
	echo ${SREADS} | xargs -n 1 > shortreads.txt
	${TIME} -v -o $(PREF).nthits.K40.time ${NTH} -b 36 -k 40 -t ${CPU} -p $(PREF).nthits --outbloom --solid @shortreads.txt
	${TIME} -v -o $(PREF).nthits.K50.time ${NTH} -b 36 -k 50 -t ${CPU} -p $(PREF).nthits --outbloom --solid @shortreads.txt
	${TIME} -v -o $(PREF).nthits.K60.time ${NTH} -b 36 -k 60 -t ${CPU} -p $(PREF).nthits --outbloom --solid @shortreads.txt
# three rounds of short-read polishing with ntEdit after two round of RACON
$(PREF).r2.nt3_edited.fa:$(PREF).r2.fa $(PREF).nthits_k60.bf
	${TIME} -v -o $(PREF).r2.nt1.time ${NTE} -t ${CPU} -k 60 -i 5 -d 5 -b $(PREF).r2.nt1 -r $(PREF).nthits_k60.bf -f $<
	${TIME} -v -o $(PREF).r2.nt2.time ${NTE} -t ${CPU} -k 50 -i 5 -d 5 -b $(PREF).r2.nt2 -r $(PREF).nthits_k50.bf -f
		$(PREF).r2.nt1_edited.fa
	${TIME} -v -o $(PREF).r2.nt3.time ${NTE} -t ${CPU} -k 40 -i 5 -d 5 -b $(PREF).r2.nt3 -r $(PREF).nthits_k40.bf -f
		$(PREF).r2.nt2_edited.fa
# all done
all: $(PREF).r2.nt3_edited.fa
```

Table S1: Short read datasets used to evaluate the performance of WENGAN.

| Sample | Technology | Machine | Read count | File | Read length | Coverage | Source |
|---|---|---|---|---|---|---|---|
| NA12878 | Illumina | HiSeq 2000 | 186,421,465 | SRR891258_1.fastq.gz | 250 | 59.97 | PRJNA196624 |
| | | | 186,421,465 | SRR891258_2.fastq.gz | | | |
| | | | 185,398,624 | SRR891259_1.fastq.gz | | | |
| | | | 185,398,624 | SRR891259_2.fastq.gz | | | |
| | Illumina | HiSeq 2500 | 266,077,618 | AH81VLADXX.R1.fastq.gz | 150 | 50.22 | GIAB |
| | | | 266,077,618 | AH81VLADXX.R2.fastq.gz | | | |
| | | | 252,850,306 | BH88WKADXX.R1.fastq.gz | | | |
| | | | 252,850,306 | BH88WKADXX.R2.fastq.gz | | | |
| | Illumina | NovaSeq | 548,283,470 | S22_L001_R1_001.fastq.gz | 150 | 53.06 | Self |
| | | | 548,283,470 | S22_L001_R1_001.fastq.gz | | | |
| | MGI | MGISEQ-2000 | 376,183,716 | EBA.bgi.fwd.fastq.gz | 150 | 36.40 | Self |
| | | | 376,183,716 | EBA.bgi.rev.fastq.gz | | | |
| | | | 172,099,754 | V100003043_L01_1.fq.gz | 150 | 16.65 | GIAB |
| | | | 172,099,754 | V100003043_L01_2.fq.gz | | | |
| NA24385 | Illumina | HiSeq 2500 | 441,957,241 | D1_S1S2_R1.fastq.gz | 250 | 71.28 | GIAB |
| | | | 441,957,241 | D1_S1S2_R2.fastq.gz | | | |
| CHM13 | Illumina | HiSeq 2500 | 202,861,861 | SRR3189742_1.fastq.gz | 250 | 66.11 | PRJNA269593 |
| | | | 202,861,861 | SRR3189742_2.fastq.gz | | | |
| | | | 206,992,396 | SRR3189741_1.fastq.gz | | | |
| | | | 206,992,396 | SRR3189741_2.fastq.gz | | | |
| HG00733 | Illumina | HiSeq 2500 | 196,489,884 | SRR5534476_1.fastq.gz | 250 | 63.26 | PRJNA300840 |
| | | | 196,489,884 | SRR5534476_2.fastq.gz | | | |
| | | | 195,745,350 | SRR5534475_1.fastq.gz | | | |
| | | | 195,745,350 | SRR5534475_2.fastq.gz | | | |
| Total | - | | 6,462,723,370 | - | - | 416.96 | - |

Table S2: Long read datasets used to evaluate the performance of WENGAN.

| | NA12878 | | CHM13 | HG00733 | NA24385 |
|---|---|---|---|---|---|
| Technology | ONT | | ONT | PacBio | ONT |
| Machine | PromethION | MinION | MinION | Sequel | MinION |
| Read count | 10,446,475 | 11,628,512 | 4,081,625 | 12,554,013 | 7,762,763 |
| Min read Size | 2,000 | 2,000 | 2,000 | 2,000 | 2,000 |
| Max read Size | 3,323,027 | 1,019,957 | 6,682,640 | 158,025 | 2,151,856 |
| N50 | 17,181 | 13,643 | 72,058 | 33,201 | 54,069 |
| N75 | 9,645 | 8,385 | 37,168 | 20,888 | 24,390 |
| Coverage | 35 | 40 | 49 | 90 | 60 |
| # reads > 100kb | 1,618 | 56,283 | 347,341 | 1,176 | 310,435 |
| Coverage reads > 100kb | 0.10 | 3.29 | 17.16 | 0.04 | 18.08 |
| Source | Self | ONT Rel5 | T2T | SRR7615963 | GIAB |

Table S3: Public long-read and hybrid assemblies of NA12878 (rel5), HG00733 (sequel), and CHM13 (T2T) used for benchmarking. All the assemblies were done by the assembler developers.

| Sample | Assembler | Version | URL | Accessed |
|---|---|---|---|---|
| NA12878 | WTDBG2 | 2.3 | NA12878.wt.fa.gz | 27/02/2019 |
| | MASURCA | 3.2.8 | MaSuRCA_3.2.8_nanopore.rel5.fa | 27/02/2019 |
| | FLYE | 2.4 | na12878.ont-ul.35x.fasta.gz | 27/03/2019 |
| | CANU | 1.7 | albacore_canu_nanopolish2_pilon2_racon2.fasta | 27/02/2019 |
| HG00733 | FALCON | Unzip v. July-2018 | RBJD01.fasta.gz | 03/05/2019 |
| CHM13 | SHASTA | 0.1 | CHM13_shasta_marginpolish_helen_consensus.fa | 14/08/2019 |
| | SHASTA | 0.1 | CHM13.shasta.fasta | 14/08/2019 |
| | CANU | 1.7.1 | chm13.draft_v0.6.fasta.gz | 28/08/2019 |
| | CANU | 1.7.1 | chm13.rel2.fasta.gz | 28/08/2019 |
| | FLYE | 2.5 | flye.v25.t2t.ont.50x.fasta.gz | 31/08/2019 |

Table S4: BAC and Fosmid sequences used to assess the consensus accuracy of genome assemblies. The sequences of NA12878 were obtained by random selecting 103 clones from an NA12878 Fosmid library. The BAC sequences of HG0073 and CHM13 were obtained from a BAC library enriched on segmental duplications.

| Sample | Sequences | Type | Total | Length (Mb) |
|---|---|---|---|---|
| NA128978 | NA12878_clones.ver_1.0.fasta | Fosmid | 103 | 3.92 |
| HG0073 | HG0073-BACs.fasta | BAC | 179 | 27.96 |
| CHM13 | CHM13-BACs.fasta | BAC | 341 | 51.53 |

Table S5: Short read assemblies. ABYSS2 and MINIA3 assemblies were run using 20 CPUs. The DISCOVAR assemblies were run using 44 CPUs and a high memory machine.

| **Sample** | **Sequencer** | **Assembler** | **Contigs** | | | **NG50** | **NG75** | **Size** | **CPU** | **Elapsed** | **RAM** |
|---|---|---|---|---|---|---|---|---|---|---|---|
| | | | **Total** | **Min** | **Max** | **(bp)** | **(bp)** | **(Mb)** | **hours** | **time** | **(Gb)** |
| NA12878 | HiSeq 2500 (2x150bp) | MINIA3 | 465,278 | 500 | 160,556 | 9,680 | 3,524 | 2,715 | 77 | 10:00:15 | 16 |
| NA12878 | NovaSeq (2x150bp) | MINIA3 | 402,298 | 500 | 189,686 | 11,797 | 4,323 | 2,722 | 69 | 8:03:09 | 15 |
| NA12878 | MGISeq (2x150bp) | MINIA3 | 428,404 | 500 | 192,702 | 10,568 | 3,739 | 2,701 | 73 | 11:01:35 | 21 |
| NA12878 | HiSeq 2500 (2x150bp) | ABYSS2 | 363,092 | 500 | 176,953 | 12,941 | 4,884 | 2,715 | 572 | 32:50:55 | 43 |
| NA12878 | NovaSeq (2x150bp) | ABYSS2 | 333,295 | 500 | 205,648 | 14,558 | 5,512 | 2,725 | 436 | 24:13:45 | 43 |
| NA12878 | MGISeq (2x150bp) | ABYSS2 | 430,581 | 500 | 192,271 | 10,789 | 3,638 | 2,682 | 437 | 24:12:22 | 43 |
| NA12878 | HiSeq 2000 (2x250bp) | DISCOVAR | 145,032 | 500 | 768,671 | 91,438 | 38,345 | 2,863 | 439 | 14:50:28 | 622 |
| NA12878 | NovaSeq (2x150bp) | DISCOVAR | 147,907 | 500 | 553,546 | 49,129 | 20,438 | 2,774 | 439 | 17:24:36 | 595 |
| NA12878 | MGISeq (2x150bp) | DISCOVAR | 157,838 | 500 | 494,557 | 43,456 | 16,964 | 2,758 | 410 | 17:33:48 | 585 |
| NA24385 | HiSeq 2500 (2x250bp) | DISCOVAR | 157,761 | 500 | 774,130 | 81,714 | 36,386 | 2,888 | 577 | 21:18:12 | 651 |
| CHM13 | HiSeq 2500 (2x250bp) | DISCOVAR | 111,955 | 500 | 823,426 | 97,044 | 42,421 | 2,853 | 723 | 23:04:00 | 647 |
| HG00733 | HiSeq 2500 (2x250bp) | DISCOVAR | 182,375 | 500 | 737,710 | 54,033 | 22,394 | 2,856 | 450 | 17:28:12 | 644 |

Table S6: QUAST validation of NA12878 assemblies (rel5). NG50 is the contig length such that using longer contigs produces half (50%) of the bases of the reference hs37d5 (3.1374 Gb) genome. NGA50 is NG50 where the lengths of the aligned blocks are counted instead of the contig lengths. LG50 is the minimum number of contigs that produces half of the reference length. LGA50 is similar to LG50 but aligned blocks are counted instead. Assembly-errors is the number of positions in the assembled contigs where the left flanking sequence aligns over 1 kbp away from the right flanking sequence on the reference (relocation) or they overlap on more than 1 kbp (relocation) or the flanking sequences align on different strands (inversion) or different chromosomes (translocation). Genome fraction (%) is the total number of bases that are aligned to the reference, divided by the reference size. The QUAST (Version: 5.0.2) analysis was run with the options min-identity 80 and fragmented ("quast -r hs37d5.fa –large –min-identity 80 –fragmented").

| Assembler | NG50 | LG50 | Assembly errors | Unaligned length | Genome fraction (%) | Largest alignment | NGA50 | LGA50 |
|---|---|---|---|---|---|---|---|---|
| WENGAND | 33,125,636 | 28 | 1,528 | 4,228,021 | 96.536 | 64,214,533 | 14,208,877 | 63 |
| WENGANA | 23,079,982 | 33 | 931 | 2,784,692 | 95.599 | 64,098,224 | 11,558,618 | 72 |
| WENGANM | 16,671,719 | 46 | 929 | 2,918,916 | 95.537 | 45,656,495 | 10,907,507 | 83 |
| FLYE | 22,178,799 | 44 | 2,598 | 11,887,494 | 96.65 | 50,090,899 | 10,557,044 | 80 |
| CANU | 10,276,017 | 81 | 2,287 | 5,590,313 | 97.051 | 33,985,615 | 6,447,156 | 129 |
| WTDBG2 | 11,672,553 | 64 | 2,030 | 25,113,146 | 92.809 | 70,481,737 | 6,294,974 | 111 |
| MASURCA | 8,243,486 | 108 | 4,379 | 8,823,738 | 97.116 | 32,622,531 | 5,209,432 | 161 |

Table S7: Fosmid evaluation using a total of 103 Fosmids (3.92Mb) of the NA12878 assemblies. "Closed" refers to the number of Fosmids for which 99.5% of their length aligns to a single locus. The identity and phred-Quality Value (QV) metrics are computed from closed Fosmids only. The common closed Fosmids are the Fosmids closed by all the evaluated genome assemblies (76 Fosmids).

| | Closed Fosmid | | Fosmid bases | | Closed Fosmid | | | | Common closed Fosmid (76) | | | |
|---|---|---|---|---|---|---|---|---|---|---|---|---|
| | | | | | Median Quality | | Mean Quality | | Median Quality | | Mean Quality | |
| | # | % | length | % | %Identity | QV | %Identity | QV | %Identity | QV | %Identity | QV |
| WENGANA | 96 | 93.20 | 3,681,736 | 93.73 | 99.83 | 27.65 | 99.43 | 22.47 | 99.86 | 28.40 | 99.62 | 24.21 |
| WENGAND | 97 | 94.17 | 3,712,540 | 94.51 | 99.92 | 30.92 | 99.55 | 23.42 | 99.93 | 31.28 | 99.82 | 27.55 |
| WENGANM | 95 | 92.23 | 3,638,627 | 92.63 | 99.80 | 27.07 | 99.53 | 23.29 | 99.83 | 27.79 | 99.68 | 24.96 |
| MASURCA | 100 | 97.09 | 3,819,651 | 97.24 | 99.80 | 26.91 | 99.69 | 25.03 | 99.80 | 27.06 | 99.74 | 25.85 |
| CANU | 94 | 91.26 | 3,584,995 | 91.27 | 99.74 | 25.88 | 99.28 | 21.45 | 99.87 | 28.78 | 99.52 | 23.23 |
| WTDBG2 | 84 | 81.55 | 3,199,569 | 81.45 | 98.06 | 17.13 | 97.93 | 16.83 | 98.06 | 17.13 | 97.96 | 16.89 |
| FLYE | 95 | 92.23 | 3,632,548 | 92.48 | 97.73 | 16.43 | 97.61 | 16.21 | 97.72 | 16.42 | 97.62 | 16.24 |

Table S8: Polishing the FLYE assembly of NA12878 with short (HiSeq 2500) and long (ONT rel5) reads. The FLYE assembly was polished using two rounds of long-read polishing with RACON followed by three rounds of short-read polishing with NTEDIT. The short-read polishing was done using the same short-reads used in the WENGAN assemblies (50X of pair-end 2x150bp reads). Consensus quality statistics after each round of polishing are presented.

| | | | FLYE | FLYE+ RACON X 1 | FLYE+ RACON X 2 | FLYE+ RACON X 2+ NTEDIT X 3 |
|---|---|---|---|---|---|---|
| T. length (Mb) | | | 2,880.84 | 2,847.89 | 2,846.09 | 2,850.65 |
| Aln. length (Mb) | | | 2,722.89 | 2,745.81 | 2,750.02 | 2,750.98 |
| bases < 99% (Mb) | | | 157.96 | 102.08 | 96.07 | 99.66 |
| Indels | short [1-2] | Number | 36,649,717 | 12,397,212 | 12,022,856 | 2,568,960 |
| | | Rate (bp) | 74 | 221 | 229 | 1,071 |
| | medium [3,50) | Number | 2,381,191 | 1,279,399 | 1,168,244 | 720,564 |
| | | Rate (bp) | 1,143 | 2,146 | 2,354 | 3,818 |
| | large >50 | Number | 13,840 | 14,544 | 14,707 | 14,903 |
| | | Rate (bp) | 196,740 | 188,793 | 186,987 | 184,593 |
| Fosmid median QV | | | 16.42 | 19.95 | 20.23 | 23.49 |
| BUSCO | | #Genes | 2268 | - | - | 3680 |
| | | %Complete | 55.3 | - | - | 89.7 |
| Computational Resources | | CPU | - | 132 | 250 | 755 |
| | | RAM | - | 386 | 386 | 386 |

Table S9: Long-read coverage tritiation of the PromethION data of NA12878 for *de novo* assembly.

| # reads | Min | Max | N50 | N75 | Bases (Mb) | Coverage |
|---|---|---|---|---|---|---|
| 1,878,641 | 5,000 | 2,914,544 | 19,575 | 12,724 | 30,000 | 10X |
| 2,817,267 | 5,000 | 2,914,544 | 19,578 | 12,728 | 45,000 | 15X |
| 3,755,076 | 5,000 | 2,914,544 | 19,590 | 12,730 | 60,000 | 20X |
| 4,694,725 | 5,000 | 3,323,027 | 19,590 | 12,729 | 75,000 | 25X |
| 5,633,658 | 5,000 | 3,323,027 | 19,589 | 12,729 | 90,000 | 30X |

Table S10: WENGAN and FLYE (v.2.5) assemblies of NA12878 at different long-read coverage. WENGAND and FLYE assemblies were run using 44 CPUs on a high memory machine (750Gb RAM). WENGANA and WENGANM were run using 20 CPUs.

| Assembler | tech | depth | length (Gb) | %Rcov 5% div | NG50 (Mb) | NGA50 (Mb) | #breaks | %Rcov 1% div | indels number short (M) | indels number medium (M) | indels number large | indels rate short | indels rate medium | indels rate large (kb) | Busco % complete | CPU hours | Max RAM (Gb) | elapsed time h:m:s |
|---|---|---|---|---|---|---|---|---|---|---|---|---|---|---|---|---|---|---|
| WENGANA | BGI | 10 | 2.738 | 87.30 | 2.91 | 2.34 | 432 | 86.05 | 2.144 | 0.467 | 16,389 | 1,259 | 5,787 | 164.73 | 93.62 | 479 | 43 | 27:16:47 |
| WENGANA | BGI | 15 | 2.748 | 87.64 | 7.70 | 4.98 | 427 | 87.08 | 1.885 | 0.412 | 16,816 | 1,450 | 6,629 | 162.47 | 94.20 | 489 | 43 | 28:05:07 |
| WENGANA | BGI | 20 | 2.753 | 87.77 | 12.19 | 6.26 | 446 | 87.34 | 1.695 | 0.386 | 16,844 | 1,617 | 7,103 | 162.69 | 94.52 | 500 | 43 | 28:51:02 |
| WENGANA | BGI | 25 | 2.755 | 87.85 | 14.19 | 6.91 | 457 | 87.43 | 1.578 | 0.370 | 16,976 | 1,739 | 7,410 | 161.59 | 94.74 | 510 | 43 | 29:37:50 |
| WENGANA | BGI | 30 | 2.757 | 87.92 | 15.46 | 6.99 | 473 | 87.52 | 1.508 | 0.362 | 17,100 | 1,821 | 7,589 | 160.58 | 94.66 | 513 | 43 | 29:46:35 |
| WENGANA | ILL | 10 | 2.750 | 87.65 | 3.63 | 2.86 | 434 | 86.76 | 1.570 | 0.419 | 17,549 | 1,733 | 6,501 | 155.12 | 94.57 | 479 | 43 | 27:13:09 |
| WENGANA | ILL | 15 | 2.757 | 87.87 | 10.40 | 6.06 | 443 | 87.39 | 1.381 | 0.380 | 17,580 | 1,986 | 7,207 | 155.97 | 94.86 | 490 | 43 | 28:06:57 |
| WENGANA | ILL | 20 | 2.759 | 87.95 | 14.19 | 6.87 | 463 | 87.54 | 1.250 | 0.362 | 17,557 | 2,198 | 7,584 | 156.44 | 94.69 | 499 | 43 | 28:59:42 |
| WENGANA | ILL | 25 | 2.761 | 88.00 | 15.81 | 7.19 | 478 | 87.64 | 1.173 | 0.352 | 17,649 | 2,345 | 7,803 | 155.80 | 94.64 | 509 | 43 | 29:39:57 |
| WENGANA | ILL | 30 | 2.762 | 88.04 | 17.02 | 7.54 | 475 | 87.68 | 1.122 | 0.346 | 17,750 | 2,452 | 7,944 | 154.98 | 94.91 | 514 | 43 | 29:52:21 |
| WENGAND | BGI | 10 | 2.772 | 88.29 | 6.84 | 4.72 | 465 | 87.52 | 1.307 | 0.338 | 16,960 | 2,101 | 8,118 | 161.91 | 94.83 | 453 | 585 | 20:16:03 |
| WENGAND | BGI | 15 | 2.777 | 88.44 | 15.39 | 7.37 | 476 | 87.93 | 1.174 | 0.308 | 17,094 | 2,350 | 8,952 | 161.38 | 94.96 | 463 | 585 | 20:53:03 |
| WENGAND | BGI | 20 | 2.780 | 88.51 | 17.16 | 7.81 | 479 | 88.09 | 1.069 | 0.294 | 17,171 | 2,585 | 9,410 | 160.95 | 94.98 | 474 | 585 | 21:36:30 |
| WENGAND | BGI | 25 | 2.781 | 88.58 | 18.03 | 7.81 | 497 | 88.16 | 1.004 | 0.286 | 17,169 | 2,755 | 9,664 | 161.10 | 95.13 | 484 | 585 | 22:11:07 |
| WENGAND | BGI | 30 | 2.783 | 88.61 | 19.40 | 7.75 | 501 | 88.20 | 0.965 | 0.283 | 17,109 | 2,867 | 9,789 | 161.74 | 95.10 | 487 | 585 | 22:20:09 |
| WENGAND | ILL | 10 | 2.776 | 88.39 | 8.04 | 5.11 | 479 | 87.71 | 1.192 | 0.324 | 18,045 | 2,308 | 8,503 | 152.50 | 95.00 | 480 | 595 | 20:05:13 |
| WENGAND | ILL | 15 | 2.780 | 88.54 | 16.37 | 7.68 | 494 | 88.04 | 1.066 | 0.298 | 18,084 | 2,590 | 9,277 | 152.74 | 94.91 | 490 | 595 | 20:39:54 |
| WENGAND | ILL | 20 | 2.783 | 88.64 | 18.03 | 8.42 | 512 | 88.19 | 0.977 | 0.286 | 18,133 | 2,832 | 9,691 | 152.58 | 94.81 | 501 | 595 | 21:25:37 |
| WENGAND | ILL | 25 | 2.785 | 88.66 | 19.40 | 8.90 | 512 | 88.23 | 0.920 | 0.280 | 18,124 | 3,009 | 9,902 | 152.74 | 94.86 | 512 | 595 | 22:03:21 |
| WENGAND | ILL | 30 | 2.786 | 88.71 | 19.40 | 8.81 | 518 | 88.28 | 0.885 | 0.276 | 18,210 | 3,128 | 10,053 | 152.10 | 94.81 | 515 | 595 | 22:14:22 |
| WENGANM | BGI | 10 | 2.756 | 87.82 | 2.53 | 2.07 | 487 | 86.80 | 1.856 | 0.426 | 17,256 | 1,467 | 6,399 | 157.81 | 94.37 | 119 | 37 | 14:09:11 |
| WENGANM | BGI | 15 | 2.765 | 88.12 | 7.09 | 4.28 | 479 | 87.63 | 1.621 | 0.378 | 17,311 | 1,696 | 7,281 | 158.81 | 95.00 | 129 | 37 | 14:55:31 |
| WENGANM | BGI | 20 | 2.768 | 88.21 | 10.06 | 5.59 | 533 | 87.83 | 1.459 | 0.354 | 17,299 | 1,889 | 7,784 | 159.29 | 94.83 | 139 | 37 | 15:47:02 |
| WENGANM | BGI | 25 | 2.770 | 88.26 | 11.35 | 5.82 | 566 | 87.91 | 1.367 | 0.342 | 17,360 | 2,018 | 8,062 | 158.88 | 94.76 | 149 | 38 | 16:31:26 |
| WENGANM | BGI | 30 | 2.772 | 88.31 | 11.92 | 6.07 | 559 | 87.94 | 1.308 | 0.335 | 17,284 | 2,110 | 8,232 | 159.63 | 94.91 | 153 | 44 | 16:41:07 |
| WENGANM | ILL | 10 | 2.758 | 87.90 | 2.89 | 2.29 | 478 | 87.01 | 1.618 | 0.418 | 17,428 | 1,687 | 6,536 | 156.63 | 94.52 | 109 | 37 | 10:57:36 |
| WENGANM | ILL | 15 | 2.766 | 88.14 | 7.40 | 4.60 | 483 | 87.68 | 1.419 | 0.375 | 17,480 | 1,939 | 7,332 | 157.38 | 94.96 | 124 | 37 | 11:58:01 |
| WENGANM | ILL | 20 | 2.770 | 88.24 | 11.07 | 5.48 | 487 | 87.83 | 1.279 | 0.356 | 17,367 | 2,154 | 7,748 | 158.66 | 94.88 | 134 | 37 | 12:52:18 |
| WENGANM | ILL | 25 | 2.771 | 88.29 | 11.68 | 5.94 | 516 | 87.90 | 1.198 | 0.346 | 17,284 | 2,302 | 7,977 | 159.55 | 94.83 | 145 | 37 | 13:41:45 |
| WENGANM | ILL | 30 | 2.774 | 88.37 | 12.26 | 5.83 | 536 | 87.96 | 1.154 | 0.340 | 17,462 | 2,392 | 8,110 | 158.04 | 94.71 | 150 | 42 | 13:47:51 |
| FLYE | ONT | 10 | 2.808 | 87.96 | 0.64 | 0.59 | 792 | 85.38 | 21.682 | 1.479 | 16,402 | 124 | 1,811 | 163.31 | 56.70 | 281 | 270 | 10:15:20 |
| FLYE | ONT | 15 | 2.807 | 88.81 | 3.75 | 2.96 | 764 | 88.01 | 13.991 | 1.072 | 17,269 | 197 | 2,575 | 159.90 | 70.88 | 370 | 334 | 14:01:46 |
| FLYE | ONT | 20 | 2.806 | 88.87 | 13.05 | 6.58 | 699 | 88.26 | 10.580 | 0.972 | 17,394 | 262 | 2,848 | 159.20 | 76.63 | 482 | 398 | 16:15:15 |
| FLYE | ONT | 25 | 2.812 | 88.92 | 16.06 | 7.22 | 822 | 88.26 | 8.912 | 0.938 | 17,341 | 311 | 2,951 | 159.69 | 80.21 | 615 | 467 | 21:04:14 |
| FLYE | ONT | 30 | 2.814 | 88.93 | 16.65 | 8.91 | 851 | 88.37 | 7.969 | 0.925 | 17,395 | 348 | 2,997 | 159.39 | 82.19 | 738 | 531 | 22:00:03 |

Table S11: Fosmid evaluation using a total of 103 Fosmids (3.92Mb) of NA12878. ”Closed” refers to the number of Fosmids for which 99.5% of their length aligns to a single locus. The identity and phred-Quality Value (QV) metrics are computed from closed Fosmids only. The common closed Fosmids are the Fosmids closed by all the evaluated genome assemblies (86 Fosmids).

| Assembler | Tech | LRC | Closed Fosmid | | Fosmid bases | | Closed Fosmid | | | | Common closed Fosmid (86) | | | |
|---|---|---|---|---|---|---|---|---|---|---|---|---|---|---|
| | | | | | | | Median Quality | | Mean Quality | | Median Quality | | Mean Quality | |
| | | | # | % | length | % | %Identity | QV | %Identity | QV | %Identity | QV | %Identity | QV |
| WENGANA | BGI | 10 | 89 | 86.41 | 3,410,705 | 86.83 | 99.77 | 26.34 | 99.58 | 23.75 | 99.77 | 26.47 | 99.59 | 23.91 |
| WENGANA | BGI | 15 | 94 | 91.26 | 3,600,050 | 91.65 | 99.82 | 27.44 | 99.39 | 22.16 | 99.83 | 27.67 | 99.65 | 24.58 |
| WENGANA | BGI | 20 | 94 | 91.26 | 3,599,919 | 91.65 | 99.83 | 27.75 | 99.68 | 24.89 | 99.84 | 27.89 | 99.67 | 24.83 |
| WENGANA | BGI | 25 | 95 | 92.23 | 3,636,440 | 92.58 | 99.83 | 27.64 | 99.71 | 25.42 | 99.83 | 27.79 | 99.72 | 25.54 |
| WENGANA | BGI | 30 | 95 | 92.23 | 3,636,440 | 92.58 | 99.83 | 27.61 | 99.72 | 25.47 | 99.84 | 28.06 | 99.72 | 25.54 |
| WENGANA | ILL | 10 | 94 | 91.26 | 3,590,963 | 91.42 | 99.86 | 28.45 | 99.69 | 25.06 | 99.86 | 28.69 | 99.74 | 25.81 |
| WENGANA | ILL | 15 | 95 | 92.23 | 3,634,145 | 92.52 | 99.87 | 28.88 | 99.75 | 26.07 | 99.88 | 29.26 | 99.76 | 26.25 |
| WENGANA | ILL | 20 | 96 | 93.20 | 3,670,535 | 93.44 | 99.87 | 28.89 | 99.76 | 26.26 | 99.88 | 29.36 | 99.77 | 26.34 |
| WENGANA | ILL | 25 | 96 | 93.20 | 3,670,535 | 93.44 | 99.87 | 28.94 | 99.78 | 26.52 | 99.89 | 29.42 | 99.78 | 26.59 |
| WENGANA | ILL | 30 | 96 | 93.20 | 3,670,535 | 93.44 | 99.86 | 28.55 | 99.76 | 26.20 | 99.88 | 29.33 | 99.76 | 26.25 |
| WENGAND | BGI | 10 | 96 | 93.20 | 3,670,535 | 93.44 | 99.89 | 29.70 | 99.76 | 26.27 | 99.90 | 29.91 | 99.78 | 26.56 |
| WENGAND | BGI | 15 | 96 | 93.20 | 3,670,535 | 93.44 | 99.90 | 29.88 | 99.77 | 26.41 | 99.90 | 29.92 | 99.78 | 26.52 |
| WENGAND | BGI | 20 | 96 | 93.20 | 3,670,535 | 93.44 | 99.90 | 29.92 | 99.79 | 26.69 | 99.90 | 30.14 | 99.79 | 26.74 |
| WENGAND | BGI | 25 | 96 | 93.20 | 3,670,535 | 93.44 | 99.90 | 29.90 | 99.79 | 26.81 | 99.91 | 30.35 | 99.79 | 26.81 |
| WENGAND | BGI | 30 | 96 | 93.20 | 3,670,535 | 93.44 | 99.90 | 30.14 | 99.79 | 26.73 | 99.91 | 30.26 | 99.79 | 26.74 |
| WENGAND | ILL | 10 | 96 | 93.20 | 3,682,021 | 93.74 | 99.89 | 29.63 | 99.72 | 25.60 | 99.90 | 30.12 | 99.77 | 26.30 |
| WENGAND | ILL | 15 | 97 | 94.17 | 3,712,392 | 94.51 | 99.90 | 30.16 | 99.75 | 26.09 | 99.92 | 30.83 | 99.78 | 26.53 |
| WENGAND | ILL | 20 | 97 | 94.17 | 3,712,392 | 94.51 | 99.91 | 30.62 | 99.76 | 26.27 | 99.92 | 31.10 | 99.78 | 26.49 |
| WENGAND | ILL | 25 | 97 | 94.17 | 3,712,392 | 94.51 | 99.91 | 30.64 | 99.78 | 26.60 | 99.93 | 31.25 | 99.79 | 26.79 |
| WENGAND | ILL | 30 | 96 | 93.20 | 3,670,535 | 93.44 | 99.91 | 30.57 | 99.79 | 26.88 | 99.92 | 30.83 | 99.80 | 26.92 |
| WENGANM | BGI | 10 | 93 | 90.29 | 3,557,496 | 90.57 | 99.85 | 28.14 | 99.64 | 24.43 | 99.87 | 28.80 | 99.68 | 24.89 |
| WENGANM | BGI | 15 | 96 | 93.20 | 3,670,535 | 93.44 | 99.85 | 28.29 | 99.64 | 24.38 | 99.86 | 28.67 | 99.65 | 24.53 |
| WENGANM | BGI | 20 | 96 | 93.20 | 3,670,535 | 93.44 | 99.86 | 28.46 | 99.68 | 24.95 | 99.87 | 29.02 | 99.68 | 24.90 |
| WENGANM | BGI | 25 | 96 | 93.20 | 3,670,535 | 93.44 | 99.88 | 29.13 | 99.72 | 25.53 | 99.88 | 29.29 | 99.72 | 25.51 |
| WENGANM | BGI | 30 | 96 | 93.20 | 3,670,535 | 93.44 | 99.88 | 29.12 | 99.73 | 25.65 | 99.89 | 29.47 | 99.72 | 25.58 |
| WENGANM | ILL | 10 | 92 | 89.32 | 3,507,285 | 89.29 | 99.87 | 28.72 | 99.73 | 25.68 | 99.87 | 29.02 | 99.74 | 25.86 |
| WENGANM | ILL | 15 | 93 | 90.29 | 3,550,467 | 90.39 | 99.88 | 29.19 | 99.75 | 25.97 | 99.88 | 29.32 | 99.75 | 26.08 |
| WENGANM | ILL | 20 | 94 | 91.26 | 3,586,857 | 91.31 | 99.87 | 28.83 | 99.76 | 26.26 | 99.89 | 29.48 | 99.77 | 26.40 |
| WENGANM | ILL | 25 | 94 | 91.26 | 3,586,857 | 91.31 | 99.87 | 29.00 | 99.77 | 26.41 | 99.89 | 29.40 | 99.78 | 26.56 |
| WENGANM | ILL | 30 | 94 | 91.26 | 3,586,857 | 91.31 | 99.87 | 28.83 | 99.77 | 26.38 | 99.89 | 29.40 | 99.78 | 26.50 |
| FLYE | ONT | 10 | 91 | 88.35 | 3,483,942 | 88.69 | 98.26 | 17.60 | 98.04 | 17.07 | 98.27 | 17.63 | 98.09 | 17.18 |
| FLYE | ONT | 15 | 96 | 93.20 | 3,676,002 | 93.58 | 98.80 | 19.19 | 98.71 | 18.88 | 98.80 | 19.20 | 98.73 | 18.96 |
| FLYE | ONT | 20 | 98 | 95.15 | 3,752,655 | 95.54 | 99.05 | 20.22 | 98.91 | 19.62 | 99.06 | 20.28 | 98.94 | 19.74 |
| FLYE | ONT | 25 | 96 | 93.20 | 3,670,535 | 93.44 | 99.14 | 20.67 | 99.03 | 20.12 | 99.16 | 20.78 | 99.05 | 20.20 |
| FLYE | ONT | 30 | 98 | 95.15 | 3,752,655 | 95.54 | 99.20 | 20.94 | 99.09 | 20.39 | 99.22 | 21.08 | 99.11 | 20.53 |

Table S12: Polishing the FLYE assembly of NA12878 with short (NovaSeq) and long (ONT flipflop) reads. The FLYE assembly was polished using two rounds of long-read polishing with RACON followed by three rounds of short-read polishing with NTEDIT. The short-read polishing was done using the same short-reads used in the WENGAN assemblies (NovaSeq 53X of pair-end 2x150bp reads). Consensus quality statistics after each round of polishing are presented.

| | | | FLYE | FLYE+ RACON X 1 | FLYE+ RACON X 2 | FLYE+ RACON X 2+ NTEDIT X 3 |
|---|---|---|---|---|---|---|
| T. length (Mb) | | | 2,814.05 | 2,806.45 | 2,806.03 | 2,817.79 |
| Aln. length (Mb) | | | 2,772.60 | 2,768.34 | 2,767.77 | 2,768.09 |
| bases < 99% (Mb) | | | 41.44 | 38.11 | 38.26 | 49.70 |
| Indels | short | Number | 7,969,035 | 11,870,633 | 12,091,389 | 1,908,590 |
| | [1-2] | Rate (bp) | 348 | 233 | 229 | 1,450 |
| | medium | Number | 925,062 | 777,251 | 776,941 | 429,712 |
| | [3,50) | Rate (bp) | 2,997 | 3,562 | 3,562 | 6,442 |
| | large | Number | 17,395 | 17,127 | 17,132 | 17,180 |
| | >50 | Rate (bp) | 159,391 | 161,636 | 161,556 | 161,123 |
| Fosmid median QV | | | 21.08 | 21.68 | 21.62 | 27.21 |
| Busco | | #Genes | 3,373 | - | - | 3,840 |
| | | %Complete | 82.19 | - | - | 93.56 |
| Computational Resources | | CPU (h) | - | 67 | 134 | 368 |
| | | RAM (Gb) | - | 270 | 270 | 270 |

Table S13: QUAST validation of the CHM13, NA24385 and HG00073 assemblies. NG50 is the contig length such that using longer contigs produces half (50%) of the bases of the reference GRCh38 (3.0882 Gb) genome. NGA50 is NG50 where the lengths of aligned blocks are counted instead of the contig lengths. LG50 is the minimum number of contigs that produce half of the reference length. LGA50 is similar to LG50 but aligned blocks are counted instead. Assembly-errors correspond to the number of positions in the assembled contigs where the left flanking sequence aligns over 1 kbp away from the right flanking sequence on the reference (relocation), or they overlap on more than 1 kbp (relocation), or else the flanking sequences align on different strands (inversion) or different chromosomes (translocation). Genome fraction (%) is the total number of bases aligned of the reference, divided by the reference size. The QUAST (Version: 5.0.2) analysis was run with the options min-identity 80 and fragmented using the autosomes plus X and Y chromosomes of GRCh38 ("quast -r GRCh38_chrom_no_alt.fa –large –min-identity 80 –fragmented").

| Sample | Assembler | NG50 | LG50 | Assembly errors | Unaligned length | Genome fraction (%) | Largest alignment | NGA50 | LGA50 | CPU hours | Max RAM | Elapsed h:m:s |
|---|---|---|---|---|---|---|---|---|---|---|---|---|
| CHM13 | FLYE-unpolished-rel2 | 57,633,823 | 19 | 5,461 | 34,966,298 | 96.341 | 75,521,408 | 25,004,847 | 39 | 3600 | 871 | - |
| | SHASTA-Polished-rel2 | 39,342,588 | 22 | 1,101 | 8,362,716 | 95.28 | 89,969,608 | 19,876,359 | 44 | - | ∼2000 | ∼24:00:00 |
| | CANU-polished-v0.6 | 70,117,841 | 16 | 6,045 | 20,877,108 | 96.752 | 75,442,419 | 25,463,052 | 38 | 219,000 | 80 | - |
| | WENGAND | 57,450,958 | 18 | 1,171 | 9,617,457 | 95.574 | 90,554,305 | 23,887,542 | 41 | 1027 | 647 | 40:07:06 |
| NA24385 | WENGAND | 50,677,160 | 19 | 1,232 | 9,724,219 | 96.195 | 75,530,184 | 25,287,000 | 42 | 910 | 650 | 47:48:55 |
| HG00073 | WENGAND | 31,108,648 | 29 | 1,032 | 8,085,299 | 95.275 | 70,705,519 | 16,930,149 | 51 | 800 | 645 | 36:20:34 |
| | FALCON | 22,334,437 | 39 | 2,410 | 15,414,934 | 96.06 | 71,678,734 | 14,607,765 | 58 | 20000 | - | - |

Table S14: BAC evaluation using a total of 341 BACs (51,5Mb) of CHM13. "Closed" refers to the number of BACs for which 99.5% of their length aligns to a single locus. The identity and phred-Quality Value (QV) metrics are computed from closed BACs only. The common closed BACs are the BACs closed by all the evaluated genome assemblies (84 BACs).

| | Closed BACs | | BAC bases | | Closed BACs | | | | Common closed BACs (84) | | | |
|---|---|---|---|---|---|---|---|---|---|---|---|---|
| | | | | | Median Quality | | Mean Quality | | Median Quality | | Mean Quality | |
| | # | % | length | % | %Identity | QV | %Identity | QV | %Identity | QV | %Identity | QV |
| WENGAND-unpolished-rel2 | 133 | 39.00 | 20,479,559 | 39.74 | 99.86 | 28.55 | 99.59 | 23.86 | 99.96 | 33.61 | 99.82 | 27.47 |
| Canu-unpolished-rel2 | 287 | 84.16 | 43,402,172 | 84.22 | 99.21 | 21.05 | 98.91 | 19.64 | 99.30 | 21.53 | 99.20 | 20.98 |
| Canu-polished-v0.6 | 280 | 82.11 | 42,501,309 | 82.48 | 99.98 | 37.04 | 99.80 | 27.05 | 99.99 | 39.57 | 99.91 | 30.63 |
| Shasta-unpolished-rel2 | 92 | 26.98 | 13,719,217 | 26.62 | 99.47 | 22.74 | 99.37 | 21.99 | 99.47 | 22.75 | 99.42 | 22.38 |
| Shasta-Polished-rel2 | 92 | 26.98 | 13,719,217 | 26.62 | 99.83 | 27.74 | 99.69 | 25.11 | 99.84 | 27.84 | 99.71 | 25.40 |
| Flye-unpolished-rel2 | 220 | 64.52 | 32,890,891 | 63.83 | 98.97 | 19.87 | 98.78 | 19.14 | 99.12 | 20.54 | 99.01 | 20.03 |

Table S15: BAC evaluation using a total of 179 BACs (27.9Mb) of HG00733. "Closed" refers to the number of BACs for which 99.5% of their length aligns to a single locus. The identity and phred-Quality Value (QV) metrics are computed from closed BACs only. The common closed BACs are the BACs closed by all the evaluated genome assemblies (41 BACs).

| | Closed BACs | | BAC bases | | Closed BACs | | | | Common closed BACs (41) | | | |
|---|---|---|---|---|---|---|---|---|---|---|---|---|
| | | | | | Median Quality | | Mean Quality | | Median Quality | | Mean Quality | |
| | # | % | length | % | %Identity | QV | %Identity | QV | %Identity | QV | %Identity | QV |
| WENGAND | 42 | 23.46 | 6,808,211 | 24.35 | 99.79 | 26.73 | 99.40 | 22.22 | 99.79 | 26.79 | 99.39 | 22.16 |
| Falcon | 80 | 44.69 | 12,313,738 | 44.04 | 99.80 | 26.89 | 99.34 | 21.80 | 99.81 | 27.12 | 99.59 | 23.83 |

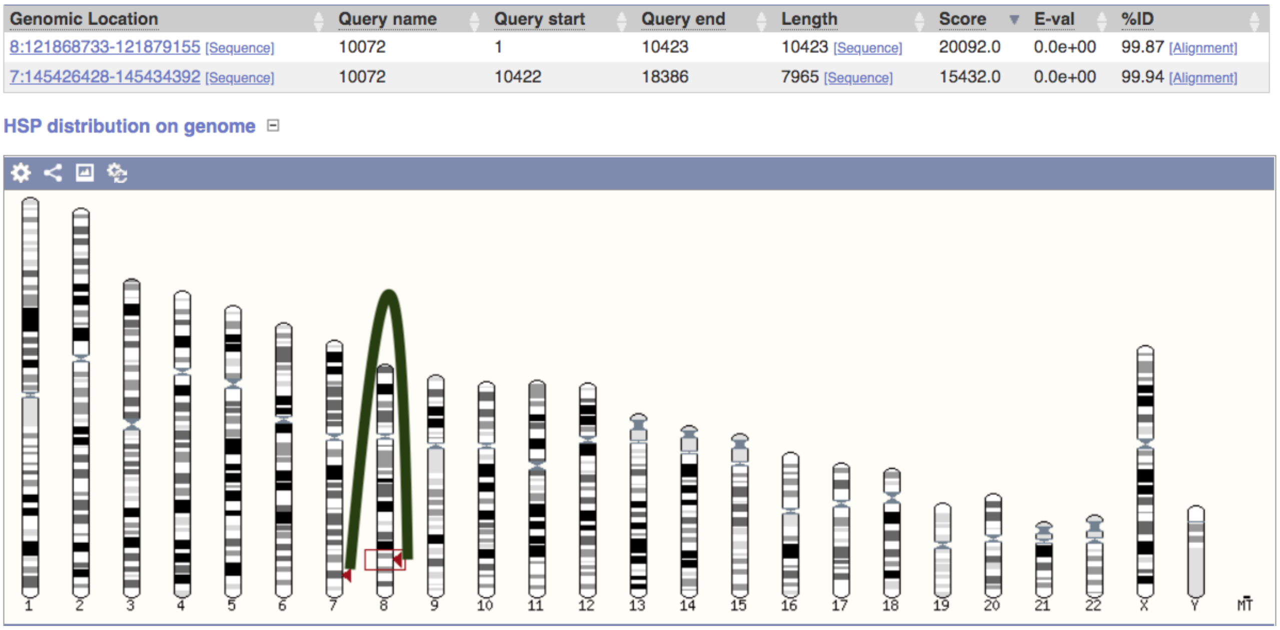

Figure S1: Example of a chimeric contig detected by IntervalMiss on the Minia3 assembly of NA12878. IntervalMiss identifies a lack of fragment coverage starting at base position 10,434 and ending at base position 10,540 of the contig 10072. A BLAST search on the human reference genome confirms the breakpoint occurring at the contig interval 10,422-10,423. In this case, the chimeric contig induces an erroneous inter-chromosome translocation between chromosomes 8 and 7. IntervalMiss splits the chimeric Minia3 contig at the flanking positions of the interval [10,433-10,540], originating in two new subcontigs covering the positions 1-10,432 and 10,541-18,386, and solving the breakpoint. The figure was generated using the Ensembl Blast portal [https://www.ensembl.org/Homo_sapiens/Tools/Blast].

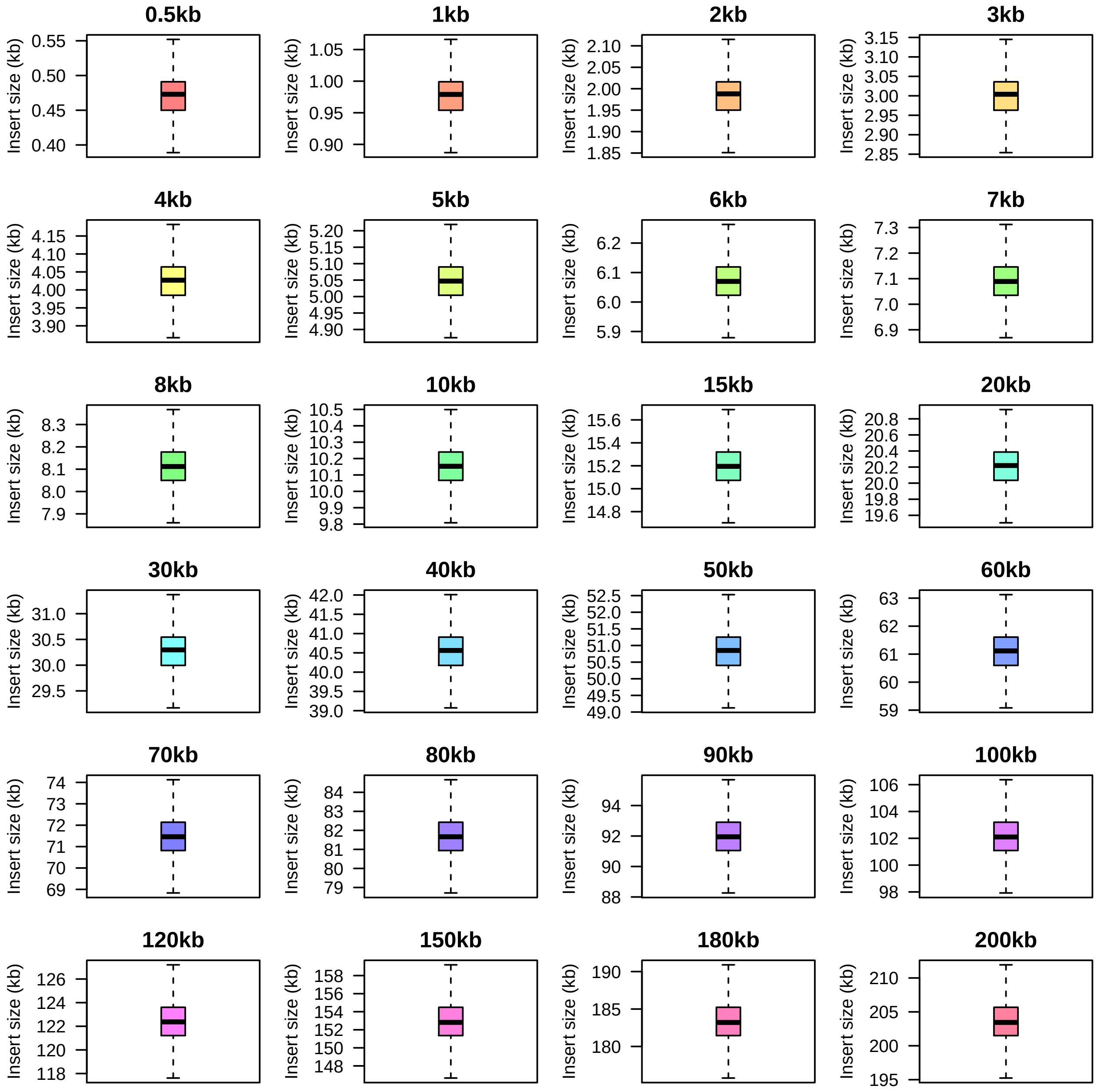

Figure S2: Spectrum of Synthetic mate-pair libraries generated by FastMin-SG from ultra-long nanopore reads of NA12878 (rel5). The boxplots were drawn extracting from the FastMin-SG alignments a minimum of 900,000 insert sizes from the mate-pair reads mapped within contigs of the NA12878 (Discovar Assembly). The percentage of outlier synthetic pairs detected ranged from a minimum of 1.18% (0.5kb) to a maximum of 16.17% (200kb).

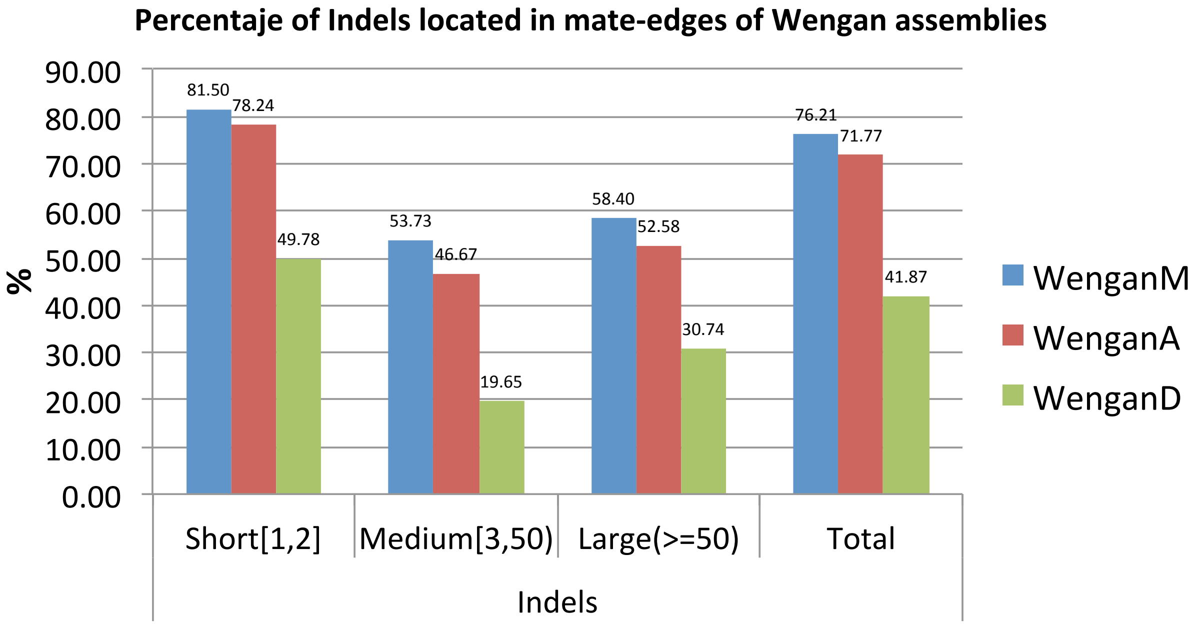

Figure S3: Distribution of the WENGAN consensus errors of the hybrid assemblies of NA12878 generated with Illumina (2x150 and 2x250) and ultra-long Oxford nanopore reads (rel5). The majority of the consensus errors are located in the long-read consensus sequences of mate-edges.

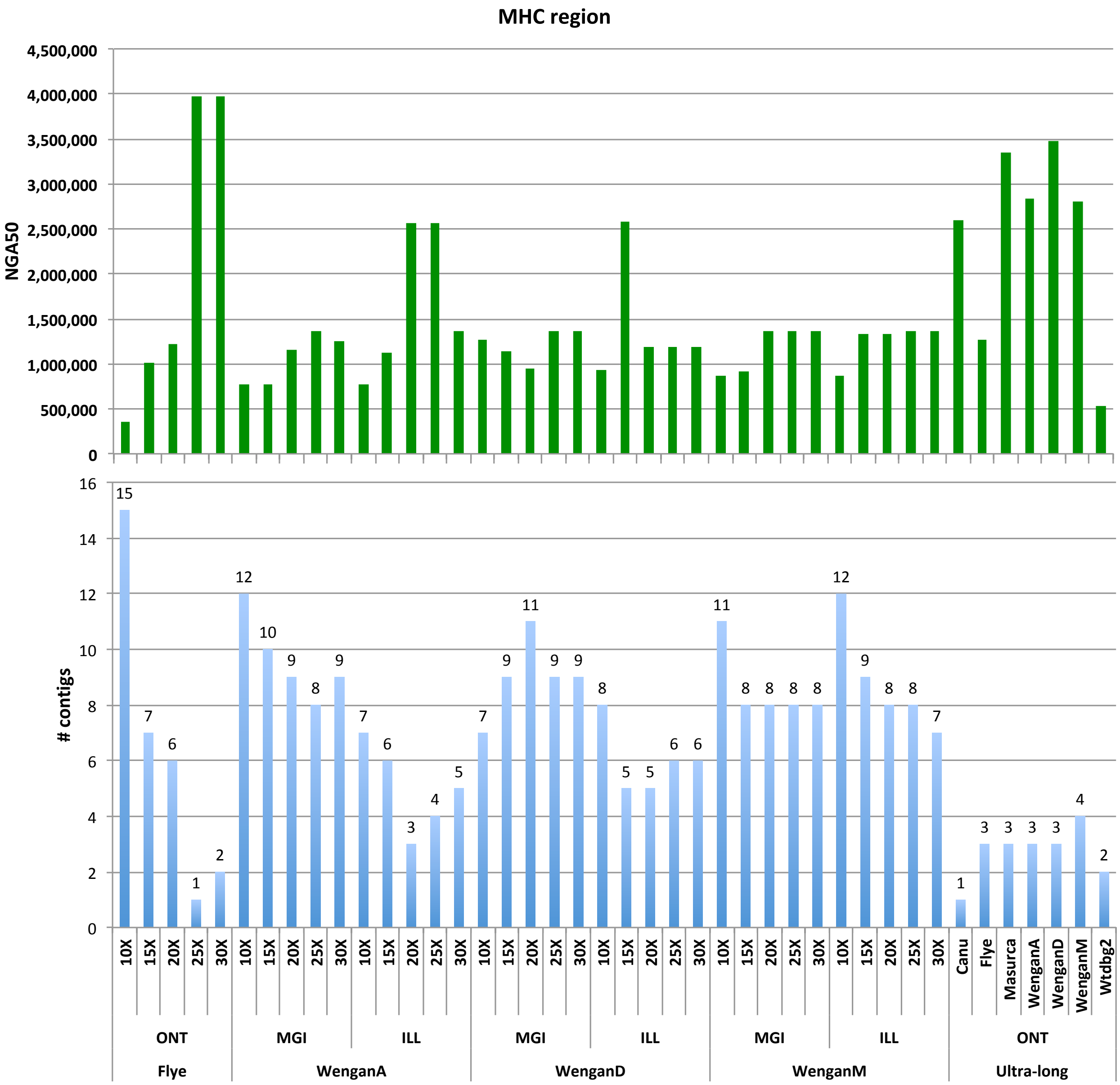

Figure S4: Assembly of the complex MHC region. The NGA50 and the number of contigs of each assembly of NA12878 are depicted. NGA50 is NG50 corrected of assembly errors.

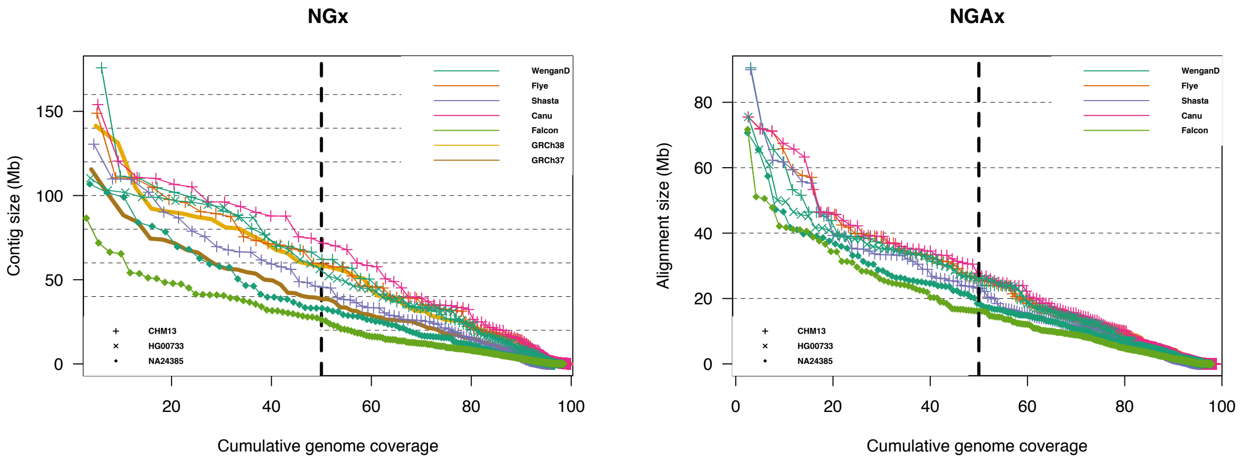

Figure S5: QUAST NGx and NGAx of HG00733, NA24385 and CHM13 assemblies.

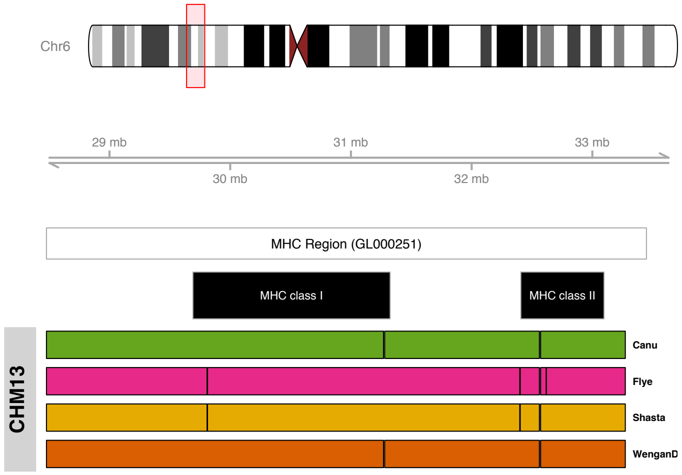

Figure S6: All the evaluated assemblers span the MHC region in a single contig. WENGAND and the curated CANU assemblies reach the higher NGA50 with a value of 2.79Mb. SHASTA and FLYE reach an NGA50 of 2.58Mb. The WENGAND contig that span the MHC region is WSC68057[1,577,866-6,402,267], with a total length of 57.4Mb. The sequence of the GL000251.2 haplogroup was used as the closed reference for CHM13.
